# Optimal Reference Panel Design in Ancient DNA Imputation from Coalescent Theory, Simulation, and Real Data Application with an Ancient Reference Panel

**DOI:** 10.64898/2026.04.27.721163

**Authors:** Bárbara Sousa da Mota, Kiran H. Kumar, David Reich, Sebastian Zöllner

## Abstract

Imputation is widely used in the ancient DNA (aDNA) field to determine which phenotypically important alleles ancient individuals carried, to study natural selection, and to detect segments of the genome that are shared between individuals identical by descent. However, rare variant imputation is less accurate, and rare variants tend to be excluded from downstream analyses. State-of-the-art imputation methods leverage large reference panels, improving rare variant accuracy in modern targets. However, it is unclear how to identify optimal panels for aDNA targets. It seems plausible that aDNA reference panels would improve imputation of aDNA, but no such panels have been assembled or tested. We leveraged analytical results from coalescent theory and complementary simulations to evaluate both performance of large modern panels, and ancient panels’ impact on aDNA imputation. For modern panels, sample sizes as small as 5,000 saturate imputation performance and model misspecifications in standard imputation algorithms increase imputation error for rare and intermediate frequency variants. For instance, for European hunter-gatherers, non-reference imputed variants with derived allele frequency less than at least 2% should be removed. Including ancient genomes in a modern reference panel substantially improved imputation accuracy in analytical modelling and simulations, particularly, for rare variants and older samples from groups with low effective population size. We assembled a joint reference panel with 1000 Genomes and 95 ancient samples and used it to impute 95 downsampled genomes, finding modest gains in imputation performance. This approach can rescue rare variants typically discarded from current imputation pipelines and may prove useful as the number of ancient samples increases.

## Introduction

Ancient DNA (aDNA) is a growing field with rapidly increasing sample size (Mallick et al. 2024; Akbari et al. 2026). However, obtaining high-quality genotypes is technically challenging, due to low yield of endogenous DNA, higher genotyping error due to deamination and potential contamination (Briggs et al. 2007; Peyrégne and Prüfer 2020). Of the more than 10,000 ancient individual samples that have been reported, only around 100 have been sequenced at high depth (Mallick et al. 2024). For the remainder, genotype imputation allows the statistical inference of diploid genotypes, provided that their sequencing results in a minimum genome coverage. The resulting genotypes facilitate subsequent population genetic inference, including identifying runs of homozygosity (Sousa da Mota et al. 2023), ancient IBD (identity by descent) segments (Ringbauer et al. 2024) and selection (Vaughn and Nielsen 2024; Irving-Pease et al. 2024; Akbari et al. 2026). Although recent ancient genomes of European ancestry have comparable imputation accuracy to modern targets, older and non-European, particularly African, samples have poorer imputation accuracy (Ausmees et al. 2022; Sousa da Mota et al. 2023).

Optimal reference panel design is critical for imputing accurate genotypes. For modern samples, key determinants include the size of the reference panel, ancestry match with the target, and minor allele frequency of variants (Das et al. 2018; Si et al. 2021). In particular, misspecification of the canonical Li and Stephens model (Li et al. 2003) affects imputation accuracy at rare variants. Model misspecification occurs because the Li and Stephen’s model implicitly assumes there is only a single best reference template (Si et al. 2021). Si et al. (2021) showed that, for modern targets and reference panels, this assumption is violated with probability ⅓ and leads to inaccurately imputed rare variants inherent to the algorithm. This error is expected to be exacerbated in ancient samples where the coalescent times with the ancient target and modern reference panel are longer than for modern targets and cannot be shortened to earlier than the age of the ancient sample by increasing reference panel size.

The most commonly used imputation reference panel for ancient DNA is the “1000 Genomes Project” dataset, consisting of 5008 phased haplotypes from 24 diverse populations around the world (Byrska-Bishop et al. 2022; Ringbauer et al. 2024; Sousa da Mota et al. 2023). Larger reference panels have also become available for aDNA imputation, including up to one million haplotypes from the UK Biobank (Rubinacci et al. 2023; Hofmeister et al. 2023; UK Biobank Whole-Genome Sequencing Consortium 2025). For modern targets, such reference panels yield dramatically better imputation results, especially for rare variants (Rubinacci et al. 2023), but it is unclear if this is the case for aDNA. While empirical assessments of aDNA imputation exist (Hui et al. 2020; Ausmees et al. 2022; Sousa da Mota et al. 2023), analytical exploration of its properties are limited (Medina-Tretmanis et al. 2024; Biddanda et al. 2022). In Medina-Tretmanis et al. (2024), the authors used coalescent-based simulations to test various phasing and imputation strategies with modern panels across a few demographic models for recovery of population structure. In Biddanda et al. (2022), a coalescent model for imputation accuracy suggests that large reference panels are not as impactful for imputing ancient targets as they are for modern ones (Huang et al. 2013; Biddanda et al. 2022; Jewett et al. 2012; Si et al. 2021).

Assembling an ancient reference panel is now possible, given the increasing number of high-quality aDNA individual samples. Furthermore, systematic work is required to explore the feasibility, optimal design, and understanding of which ancient samples would benefit most from the implementation of such a panel. Imputation of ancient genomes may benefit substantially from a panel that contains ancient haplotypes from comparable ancestries or time periods. Building such a panel would require a significant bioinformatical effort, including calling *de novo* mutations and handling potential phasing errors.

Here we study how including ancient samples in a reference panel affects DNA imputation as well as other aspects of reference panel design by combining analytical models, simulations and real aDNA data. In our analytical models, we explore the relationship between the target haplotype and the most closely related template haplotype in a coalescent framework following (Biddanda et al. 2022). The expected coalescent time between the target and the closest template haplotype serves as a proxy for imputation error, and we can calculate it for different parameter settings. We contrast these analytical predictions with simulations with recombination using parameters from (Kamm et al. 2019). Combining analytical modelling and simulations allows us both to describe the impact of reference panel choice in a general way and focus on complex, realistic demographic scenarios. In both analyses, we first assess the impact of large, modern day reference panels on aDNA imputation; specifically, the effect of reference panel size in a structured population and rare variant imputation accuracy. The benefit of large reference panels is limited as sample age increases. We then investigated the potential benefit of adding aDNA haplotypes to a modern reference panel. We computed the analytical and simulated gain in imputation accuracy when adding 5 - 500 ancient samples into a modern reference panel, showing that, in some scenarios, even the addition of a handful of aDNA samples substantially improves imputation accuracy. By assembling a reference panel containing the 1000 Genomes panel (Byrska-Bishop et al. 2022) and 95 ancient genomes, we show that, in practice, this gain in accuracy is limited by data quality and sample size.

## Results

### Modelling framework

We perform both analytical calculations and simulations to assess aDNA imputation under different reference panels. In our analytical calculations, we consider the effect of imputing a single target variant with a reference panel consisting of *K* phased haplotypes, where 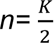 is the number of individuals. We focus on biallelic sites for simplicity, where the ancestral allele is coded as 0 and the derived allele is coded as 1. We assume that the imputation algorithm correctly selects the most closely related template haplotype. We model the relationship between those two haplotypes in a coalescent tree. The target aDNA with age *T_a_* first coalesces with a reference haplotype at time *T_c_* (**Figure 1a**). *E[T_c_]* is our measure of expected imputation error for aDNA targets, since mutations impacting imputation accuracy occur on a branch of this length (**Supplementary Note 1)**.

**Figure 1:**
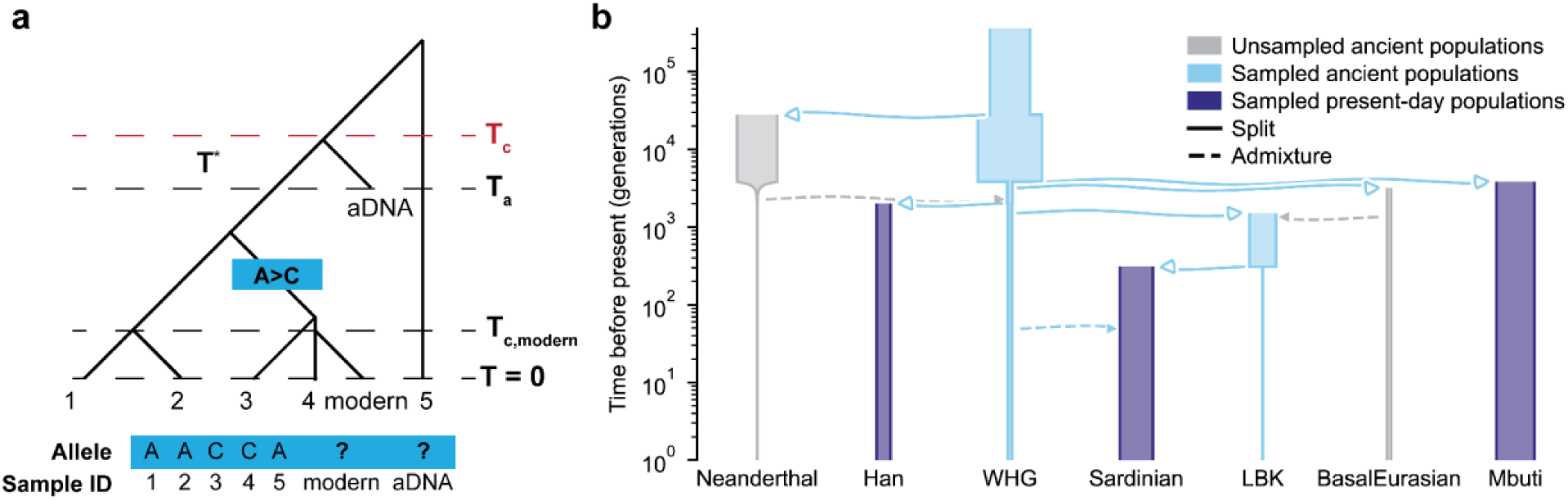
Overview of Analytical and Simulation Models. **a.** An example coalescent tree for coalescent modelling of aDNA Imputation at a single locus: here, we have five reference panel haplotypes (1 - 5) and one aDNA and one modern target. The alleles on these five haplotypes are generated from the tree structure. A is the ancestral allele, which is carried by reference panel haplotypes 1, 2, and 5. The A→ C mutation occurs on the ancestor of haplotypes 3 and 4; therefore, haplotypes 3 and 4 carry the derived allele C. The age of the aDNA target is T_a_, which is known, and the aDNA target’s first time of coalescence with the modern-day reference panel is T_c_. T* is the difference between T_c_ and T_a_. The best template(s) are those with the shortest E[T_c_] with the target, so for the modern target they are reference haplotype 4 and for the ancient target they are reference haplotypes 1-4. **b.** The demographic model at the basis of our coalescent-based simulations. Parameters are as in (Kamm et al. 2019). Bar widths scale with effective population sizes.

In our simulations of genetic data, we follow a semi-realistic population history model (Kamm et al. 2019) **(Figure 1b)** using msprime (Baumdicker et al. 2022). To mimic a subset of the 1000 Genomes panel (Byrska-Bishop et al. 2022), we sampled genomes from three populations at present time (t=0), representing African (‘Mbuti’), East Asian (‘Han’) and European (‘Sardinian’) continental groups. For ancient genomes (t>0), we have two versions of the simulations: 1) sampling of individuals from the ‘Sardinian branch’ at different time points, and 2) contemporaneous sampling WHG-like (Western Hunter-gatherer) and LBK-like (Linear Pottery culture) individuals, that have low and high effective population sizes, respectively (**Supplementary Note 2**). For both analytical and simulation models, we assume that there are no genotyping and phasing errors for the reference haplotypes or aDNA damage. In addition, for the analytical model, we assume no genotyping or phasing errors in the targets.

### Imputing aDNA with Modern Reference Panels

First, we assessed aDNA imputation performance using modern reference panels, starting by evaluating the impact of reference panel size. In our analytical modelling, we calculated the expected coalescence time between target and templates (E[T_c_]), a proxy for expected imputation error, as a function of aDNA sample age, T_a_ (years before present) (**Methods**). We assessed E[T_c_] for panel sizes of 100, 500, 1000, and 10,000 under both the unstructured and structured coalescent with three demes and a low migration rate between them (**Methods**).

Our simulations have similar parameters to our analytical model. We imputed simulated 1x genomes from ancient individuals sampled at different time points in the past from the ‘Sardinian’ lineage using a reference panel consisting of three populations with no migration between them: ‘Han’, ‘Mbuti’, and ‘Sardinian’. We varied the number of present-day ‘Sardinian’ samples while fixing the number of ‘Mbuti’ and ‘Han’ samples to 500 each. We calculated the non-reference discordance (NRD) for modern reference panels containing 100, 500, 5000, and 10,000 ‘Sardinians’, yielding total panel sizes of 1100, 1500, 6000, and 11,000 individuals.

For the analytical modelling, across reference panel sizes, the expected coalescence time between the ancient target and reference haplotypes (i.e. E[T_c_]) increased with increasing aDNA target age regardless of the existence of spatial structure (**Figure 2a**). While adding more reference haplotypes improved imputation performance, imputation accuracy saturated quickly. Modern panel sizes between 5,000 and 10,000 individuals led to approximately the same E[T_c_], for sample ages older than ∼250 years. (**Figure 2a**). Increasing panel size from n=100 to n=500 samples benefitted younger aDNA targets: for targets ∼2000 years old, adding 500 samples compared to 100 lowered E[T_c_] from 0.02 to 0.01. This benefit disappeared for older samples. For instance, at T_a_ ∼ 20,000 years old, E[T_c_] is essentially the same for n=100 and n=500. The presence of spatial structure in the reference panel and its particular parameterization only impacted our results when the number of demes was large and migration rates were low (**Supplementary Fig. 1**). Furthermore, when we calculate E[T_c_] under instantaneous population growth, the expected imputation error across all panel sizes was essentially unchanged except for small differences in the smallest panel size (n=100) and older targets (> 10,000 years before present) (**Supplementary Fig. 2).**

**Figure 2:**
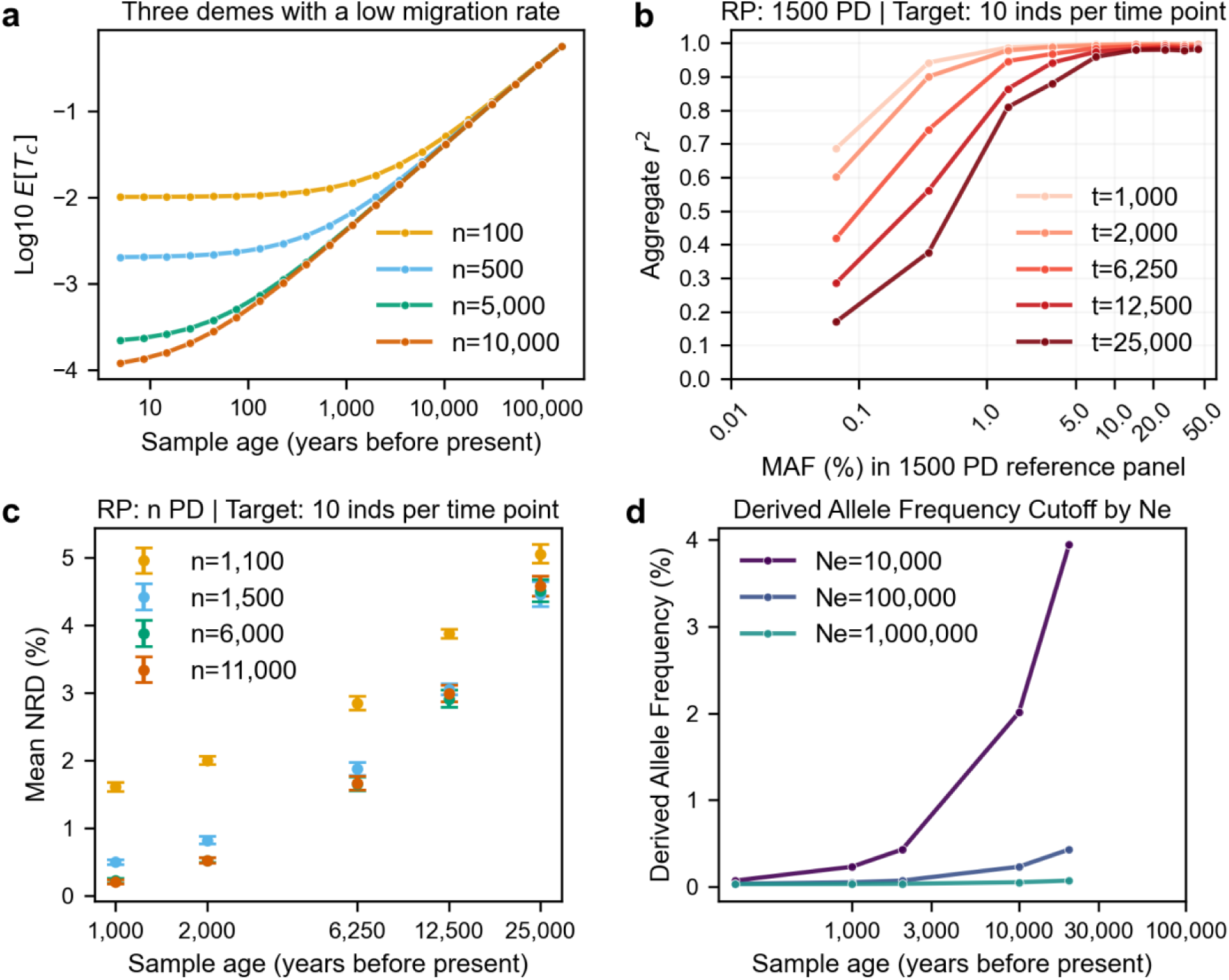
Modern reference panels have limited utility for aDNA imputation. **a.** Expected coalescence time between modern templates and aDNA target as a function of target age in units of years (assuming N_e_=10,000 individuals), with both axes in log scale. Each line denotes a different modern reference panel size. **b.** Squared Pearson correlation (r^2^) between imputed dosages and ground truth as a function of minor allele frequency (MAF) when ancient genomes of different ages were imputed with a reference panel containing 1,500 present-day (PD) genomes (500 ‘Sardinians’). **c.** Mean non-reference discordance (NRD) resulting from imputing aDNA as a function of sample age with modern reference panels of varying sizes. We varied the numbers of ‘Sardinian’ genomes, i.e. the most genetically similar reference population, while fixing the ‘Han’ and ‘Mbuti’ sample sizes to 500 each. Each data point corresponds to the mean NRD calculated for ten individuals of the same age and error bars represent two times the standard error. **d.** Suggested derived allele frequency (DAF) cutoff (%) for variants imputed as non-reference by sample age (years before present). Since the DAF is roughly equal across reference panel sizes, we specify a panel size of n=3,000 in our calculations. Each line denotes a different N_e_ (effective population size). In this figure, generation time was assumed to be 25 years. RP: reference panel.

In simulations, the target age effect on imputation performance was most severe for rare variants (MAF<2%, **Figure 2b**). For MAF between 0.1% and 1% and a reference panel containing 500 ‘Sardinians’, aggregate r^2^ is twice as high for the youngest samples (1,000 ybp, years before present) as for the oldest samples (25,000 ybp). Adding 500 ‘Sardinians’ to the reference panel compared to 100 led to a decrease in NRD from 1.60% to 0.50% at t = 1,000 ybp (**Figure 2c**). However, for older targets, changing the panel size from n=100 to n=500, led to a much smaller benefit. For instance, at t = 25,000 ybp NRD reduced marginally from 5.06% to 4.46%. Increasing the reference panel size had the most impact on imputed rare variants with substantial benefits for samples younger than 6,250 ypb (**Supplementary Fig. 9** and **Supplementary Fig. 10**). The gain in imputation performance decreased with target age and was systematically smaller as the reference panel size increased. However, for 25,000 ybp, adding extra reference haplotypes was detrimental to imputation of rare variants (MAF between 0.1% and 1%). As in the analytical modelling, improvements in imputation accuracy stagnated when the reference panel size was between 5,000 and 10,000, even for younger samples (**Supplementary Fig. 9** and **Supplementary Fig. 10**), including t = 1,000 ybp for whom NRD only decreased from 0.22% to 0.20% (**Figure 2c**).

We next assessed the effect of Li and Stephens model misspecification on aDNA imputation accuracy. The Li and Stephens model implies there is only one best template from the reference panel. If this assumption is wrong, templates with younger variants may add noise to the imputation estimate. We calculated the probability distribution of the number of best templates for reference panels of sizes n = 3,000 and n = 100,000, corresponding to the sizes of 1000 Genomes and TopMed, respectively, and for aDNA samples from T_a_ = 2,000 to T_a_ = 50,000 years old. For the analytical model, the Li and Stephens model is misspecified at T_a_ = 2,000 years and n=3,000 individuals with probability of 0.95 (**Supplementary Table 1**). This is much greater than the misspecification probability of ⅓ for modern targets, regardless of the reference panel size (Si et al. 2021). For older aDNA targets (T_a_ = 20,000 years old and T_a_ = 50,000 years old) and larger reference panels (n = 100,000), this model misspecification occurs with a probability of approximately 1 (**Supplementary Fig. 3; Supplementary Table 1**).

We then calculated the expected number of templates (**Table 1**) used in the Li and Stephens model. This expectation provides a derived allele frequency cutoff (DAF) at which variants that are imputed with the derived allele are potentially younger than the coalescence time between the ancient sample and the modern reference panels templates (i.e. T_c_), leading to imputation errors inherent in the Li and Stephens model. Across sample ages, this DAF cutoff is roughly equal regardless of panel size, though the derived allele count varies substantially. For N_e_ =10,000 individuals and samples that are T_a_ = 10,000 years old, the DAF cutoff for imputed variants should be around 2% to remove mutations accrued more recently than the coalescence time between target and templates.

**Table 1:**
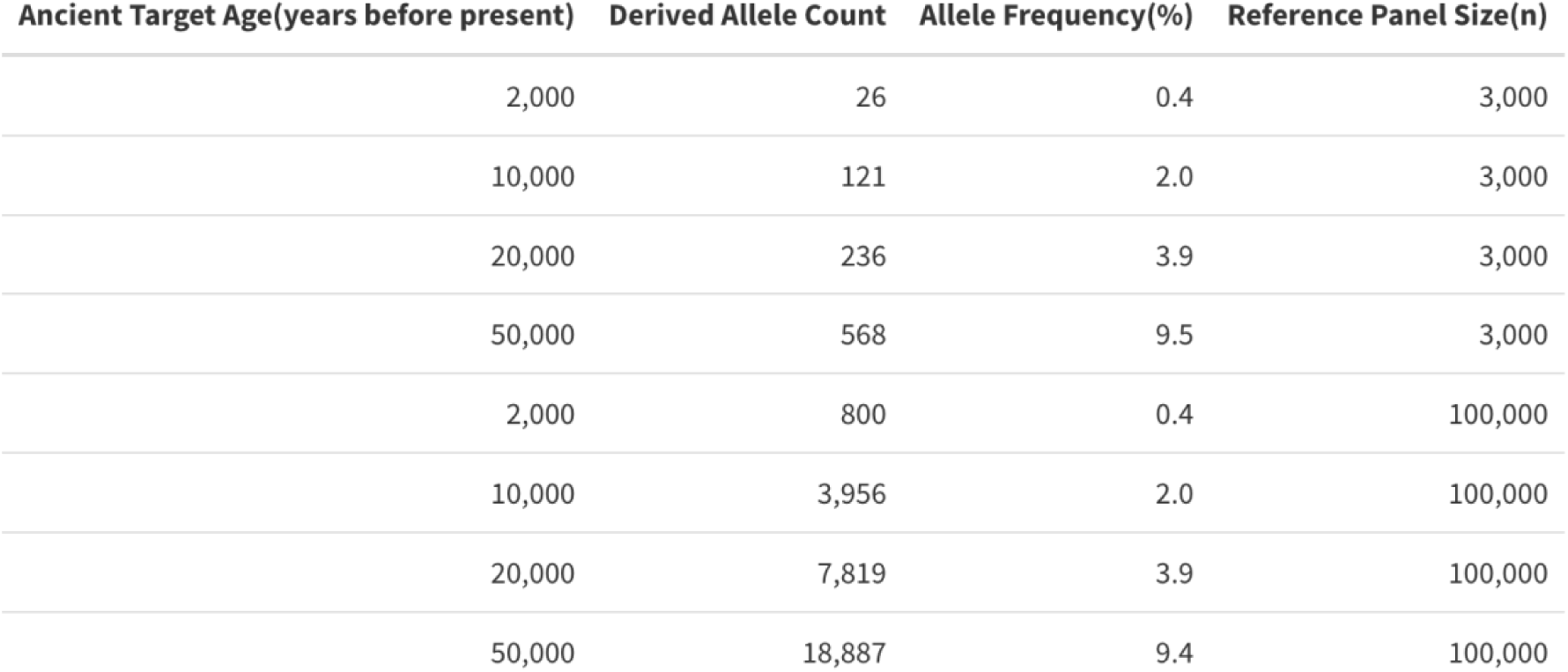
Allele frequency cutoffs for variants incorrectly imputed as non-reference due to model misspecification by age (years before present) when effective population size is 10,000 individuals.

| Ancient Target Age(years before present) | Derived Allele Count | Allele Frequency(%) | Reference Panel Size(n) |
| --- | --- | --- | --- |
| 2,000 | 26 | 0.4 | 3,000 |
| 10,000 | 121 | 2.0 | 3,000 |
| 20,000 | 236 | 3.9 | 3,000 |
| 50,000 | 568 | 9.5 | 3,000 |
| 2,000 | 800 | 0.4 | 100,000 |
| 10,000 | 3,956 | 2.0 | 100,000 |
| 20,000 | 7,819 | 3.9 | 100,000 |
| 50,000 | 18,887 | 9.4 | 100,000 |

We note that the relationship between DAF and sample age is linear. For instance, for N_e_ = 10,000 individuals and samples that are 2,000 years old the DAF cutoff is 0.4%, which is a five-fold decrease in both age and DAF cutoff. This linear relationship implies that samples that are 5,000 years old require a DAF cutoff that includes intermediate frequency variants (DAF > 1%). For a given sample age, as N_e_ increases, the derived allele frequency cutoff decreases (**Figure 2d**). For instance, for 20,000 ybp, when N_e_ = 100,000 individuals the cutoff is 0.4% and when N_e_ = 1 million individuals, the cutoff is 0.07%, which is considerably smaller than the 3.9% cutoff for N_e_ = 10,000 individuals.

### Adding Ancient Samples to a Modern Reference Panel

Both coalescent analytical modelling and simulation indicate that adding ancient samples to a modern reference panel substantially increases imputation accuracy. We assessed imputation error while adding 0 - 500 aDNA samples of the same age as the target to a modern reference panel of size n= 1,500.

To test the impact of distinct demographic histories, we added aDNA to a modern reference panel under different constant, fast growth and ‘low’ and ‘high’ N_e_ scenarios. For the analytical derivations, we modelled two different demographic histories, one with a constant Ne of 10,000 individuals and the other with instantaneous population growth from N_e_=10,000 individuals to N_e_ = 10 million individuals at t = 0.03 2N_e_ generations, which is chosen to match the harmonic mean of N_e_ under exponential growth for Europeans (Gutenkunst et al. 2009) (**Supplementary Note 1**). Correspondingly, we simulated ancient individuals from two different populations, namely, European hunter-gatherer-like (‘WHG’) and LBK-like with ‘low’ and ‘high’ effective population sizes of 1,920 and 12,000, respectively.

Regardless of N_e_, as the aDNA target age increased, the expected imputation error from the analytical modelling decreased (**Figure 3a-b**). Across all aDNA ages, the reference panel with no ancient samples has the largest expected imputation error (i.e. E[T_c_]). For the analytical model with constant population size, for targets aged ∼30,000 years old, adding ten ancient samples decreased E[T_c_] from 0.12 to 0.075 and adding 100 ancient samples further decreased it to 0.02. Adding up to 500 ancient samples did not saturate E[T_c_], which reduced to 0.004 **(Figure 3a)**. However, for the analytical model with fast growth from N_e_ = 10,000 individuals to N_e_ = 10 million individuals, adding 10 ancient genomes provided virtually no benefit and even for adding 500 ancient genomes the expected imputation error only decreased from 0.21 to 0.16 in ∼10,000-year-old targets. For targets that are ∼18,000 years old, which is just before the instantaneous growth, adding 500 ancient samples lowered E[T_c_] from 0.04 to 0.01 **(Figure 3b)**.

**Figure 3:**
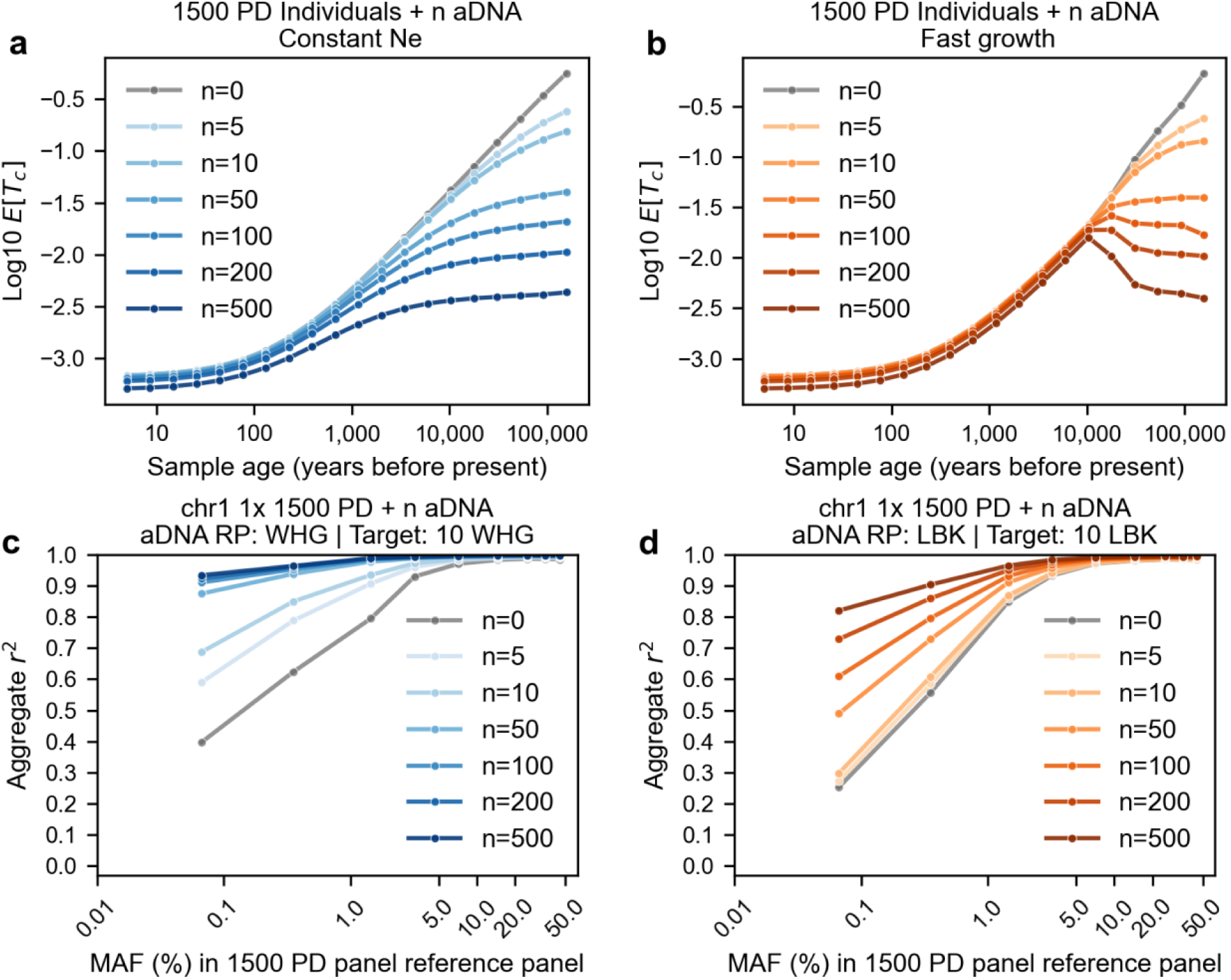
Impact of adding aDNA haplotypes to a modern reference panel under different demographic models. **a-b** The expected coalescence time between target and best templates for aDNA imputation with 1,500 present day (PD) individuals and between 0 to 500 ancient samples in a reference panel across aDNA target ages (in units of years before present). We display this under constant low N_e_=10,000 **(a)** and high N_e_ **(b)** after instantaneous population growth from N_e_=10,000 individuals to N_e_=10 million individuals at time = 0.03 2N_e_ generations. We assume that the generation time is 25 years. **c-d** Imputation accuracy (aggregate r^2^) as a function of minor allele frequency (MAF) when we vary the number of ancient genomes in the imputation reference panel between 0 and 500. **c.** Imputation of ten WHG-like genomes while adding other ‘WHG’ haplotypes to a reference panel of 1,500 PD genomes. **d.** Imputation of ten ‘LBK’ genomes using a reference panel containing other ‘LBK’ genomes. WHG: Western hunter-gatherers; LBK: Linear Pottery Culture; aDNA RP: aDNA reference panel.

In simulations, an imputation panel without aDNA led to the smallest aggregated r^2^, while the reference panel with 500 ancient samples led to the largest aggregated r^2^. These improvements were particularly large for older ancient individual samples and rare variants (**Supplementary Fig. 11** and **Supplementary Fig. 12**). For instance, when imputing 25,000-year-old samples, adding 50 aDNA genomes from 12,500 years before present to the reference panel increased r ^2^ at MAF<0.1% from 0.17 to 0.46, and adding 500 aDNA genomes in total led to r^2^=0.69 (**Supplementary Fig. 12**). Furthermore, imputation error rates decreased regardless of the mutation age (**Supplementary Fig. 13**).

In simulations of WHG-like and LBK-like groups, imputation accuracy for ancient target genomes increased with increasing number of aDNA reference haplotypes when the ancestries were well matched. In other words, reference haplotypes from ancient populations with a rather distinct population history from the target genomes, such as the ‘WHG’ and ‘LBK’ in our simulations, have little to no effect on imputation performance (**Supplementary Fig. 14**). For the low-N_e_ scenarios, including as few as five ancient genomes in the reference panel led to marked improvements. For the simulations with the ‘WHG’ genomes, r^2^ increased from 0.40 to 0.59 for variants in the lowest MAF bin (0-0.1%), and r^2^ saturated at ∼0.90 for n=100 reference genomes (**Figure 3c**). When imputing LBK-like genomes, a minimum of 50 reference genomes was required for imputation performance to substantially improve (for MAF<0.1%, r^2^=0.25 and r^2^=0.49 for n=0 and n=50 ‘LBK’, respectively) and even at n=500 r^2^ values had not saturated (r^2^=0.82 for MAF<0.1%, **Figure 3d**). We observed similar trends when we increased N_e_ for ‘LBK’ to a maximum of 100,000, albeit with smaller imputation accuracy gains at rare variants (**Supplementary Fig. 15** and **Supplementary Fig. 16**).

Overall, for both analytical and simulated models, we observed gains in imputation accuracy when adding ancient haplotypes to modern reference panels. We saw greater imputation accuracy gains for ancient individuals from populations with smaller N_e_.

### Real Data: Imputing with aDNA samples + 1000 Genomes

In our analytical modelling and simulations, we investigated imputation accuracy of ancient-like genomes in a perfect data framework. In reality, aDNA reference panels face several challenges: postmortem damage in DNA can increase genotype calling errors that can be propagated to target samples through imputation. Moreover, phasing errors in the aDNA panel can limit potential imputation gains. As such, we examined the potential of aDNA imputation panels by assembling a dataset of 95 high coverage ancient human genomes, unrelated to the 2nd degree, from the past 45,000 years (**Figure 4a**, **Supplementary Table 7**, and **Supplementary Fig. 17**). To minimize genotype calling errors, we only called genotypes at the 1000 Genomes (1KG) biallelic SNPs on chromosome 1 and subsequently imputed these before merging with 1KG. We verified that using imputation to refine genotype calls for aDNA can lead to fewer genotype errors (**Supplementary Note 4**) and improved phasing (**Supplementary Note 5**). We then assessed imputation accuracy using a reference panel containing aDNA haplotypes by downsampling the ancient genomes to 1x and imputing following a leave-one-out approach. Imputation errors were either approximately equal or lower when using the panel with aDNA (1KG_aDNA) compared to 1KG alone (**Figure 4b**). As with the simulations, imputation accuracy increased the most for rare variants (**Figure 4c**). This was particularly the case for individuals from the European Mesolithic, from which our dataset has 20 in total, for instance, r^2^ gains varied between 0.04 and 0.15 for the rarest variants with MAF<0.1% (**Figure 4c,d**).

**Figure 4:**
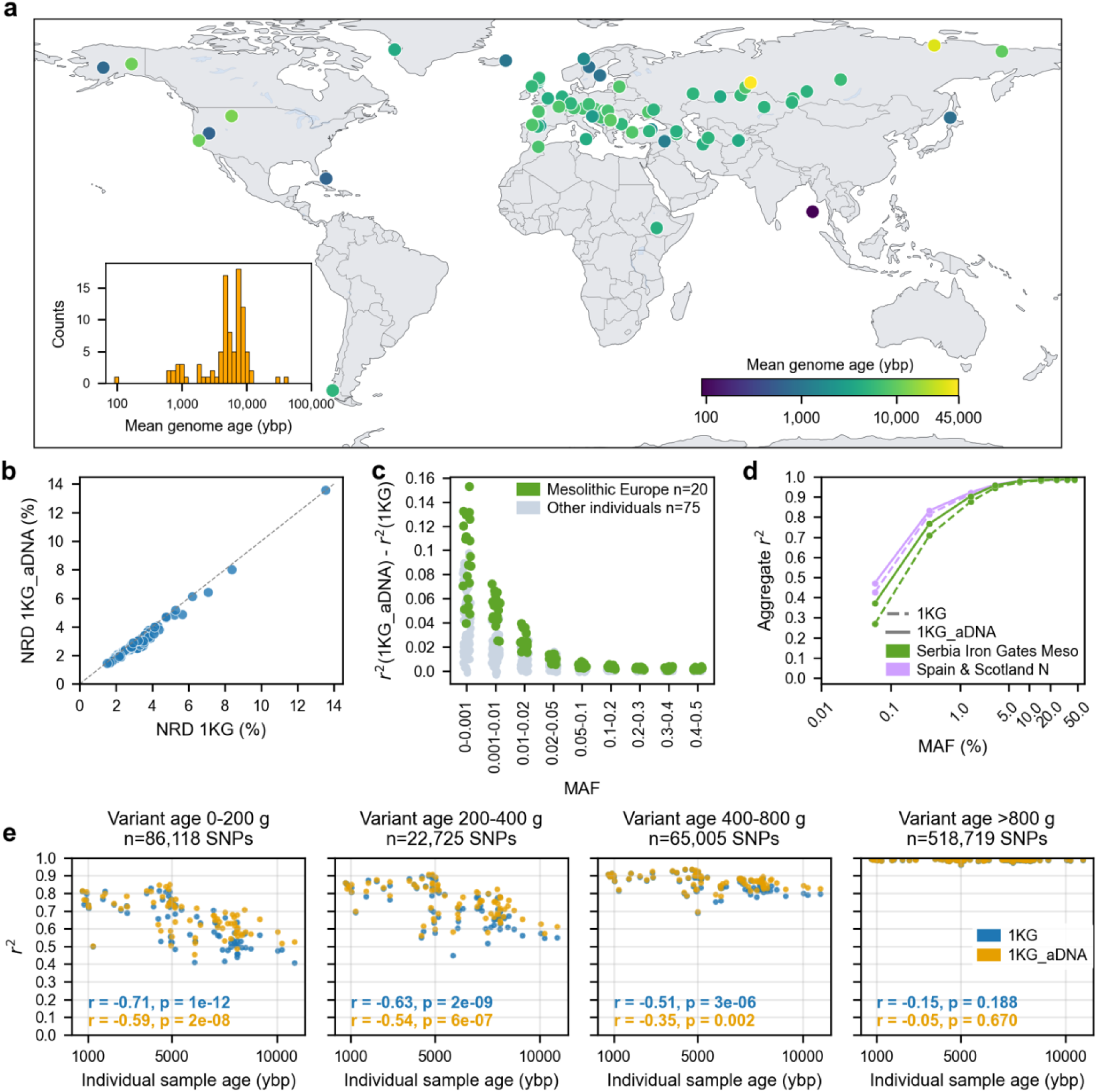
Imputation reference panel containing present-day and ancient haplotypes. **a.** Spatiotemporal distribution of the 95 high coverage ancient genomes. **b.** Comparison of NRD (non-reference discordance) values after imputing downsampled (1x) versions for each of 95 ancient genomes in a leave-one-out approach when using the 1000 Genomes reference panel (1KG) and a merged version of the 1000 Genomes and the ancient haplotypes (1KG_aDNA). **c.** Difference in aggregate r^2^ obtained using the 1KG_aDNA and 1KG panels as a function of minor allele frequency (MAF) bin for each individual sample; aggregate r^2^ is the squared Pearson correlation between imputed dosages and high coverage genotypes in a particular MAF bin. **d.** Aggregate r^2^ for two groups of individual samples, Serbia Iron Gates Mesolithic (n=11) and Spain & Scotland Neolithic (n=6), when distinct imputation reference panels were used. **e.** The four plots depict r^2^ per individual sample from West Eurasia as a function of the mean sample age across SNPs with different ages as estimated with Relate for Northern Europeans in the 1000 Genomes panel (CEU). The data points in blue and yellow indicate results obtained when using 1KG and 1KG_aDNA, respectively. Pearson correlation (r) between r^2^ and sample age for the two imputation experiments and respective p-values (p) are shown on the plots at the bottom.

We further examined how variant age and sample age impact imputation performance. To this end, we calculated imputation r^2^ for every West Eurasian individual from the past 11,000 years and for variants binned by age, as estimated by Relate on CEU (Utah residents with Northern and Western European ancestry) individuals from 1000 Genomes (Speidel et al. 2019). We found significantly negative Pearson correlations (r) between imputation r^2^ and sample age for variants with estimated age below 800 generations and the correlation magnitude decreased with variant age (**Figure 4e**). We verified that this relationship between sample age and imputation accuracy is indeed driven by variant age rather than allele frequency (**Supplementary Fig. 22**), despite these two being strongly correlated. For most individual samples, imputation accuracy was higher when using the 1KG_aDNA panel, which was more evident for younger variants and older samples. That is further illustrated by systematically lower Pearson correlation between imputation r^2^ and sample age across all variant age bins. For example, for variants younger than 200 generations, r=-0.71 and r=-0.59 for 1KG and 1KG_aDNA, respectively.

## Discussion

We provide a comprehensive assessment of aDNA imputation accuracy using both modern and ancient reference panels through analytic modelling, simulation, and application to real data. Unlike for modern targets, large reference panels do not necessarily facilitate better accuracy, especially for older samples. Furthermore, modern reference panels may incorrectly impute derived alleles younger than the coalescence time of the target and the reference haplotypes due to model misspecification of the Li and Stephens algorithm. The frequency of such variants depends strongly on sample age. Adding aDNA haplotypes to the reference panel can, however, substantially improve imputation performance of ancient genomes. We evaluated this prediction by assembling a dataset of 95 high-quality ancient genomes and merged it with the 1000 Genomes panel and assessed imputation performance. We achieved modest gains in imputation accuracy that were most prominent for European hunter-gatherers.

Our results have important implications for designing aDNA imputation studies. In practice, many studies discard variants with low MAF due to low imputation accuracy (e.g., (Allentoft et al. 2024; Cassidy et al. 2025; Akbari et al. 2026)). However, rare variants constitute the vast majority of human variation, are important in understanding fine population structure (Huang et al. 2025), and tend to have larger functional impacts on phenotypes (Wang et al. 2021; Karczewski et al. 2022; Sella and Barton 2019). Our results indicate that discarding variants with MAF < 1% may be too liberal for older samples (e.g. European hunter-gatherers) and a more appropriate cutoff for them would be a derived allele frequency of at least around 2%. Furthermore, imputation accuracy for aDNA saturates at around 5,000 modern individuals in the reference panel.

In simulations, we observed the largest gains in imputation accuracy at rare variants when we added aDNA to a modern reference panel. This was particularly the case for targets from low N_e_ populations, for which we found that as few as five aDNA reference haplotypes considerably boosted imputation performance. Importantly, matching the ancestry of the target population with the ancient reference haplotypes was key to this improvement. However, aDNA imputation references with unmatched ancestries did not decrease imputation accuracy in both simulations and real data.

Compared to our analytical and simulated predictions, increases in imputation accuracy for adding aDNA references in real data were smaller, likely due to the diverse spatiotemporal distribution of the aDNA samples and the well-documented data quality concerns with aDNA, particularly with genotyping and phasing errors. We assumed no such errors in our analytical modelling and simulations, and, in particular, in the analytical modelling, we assumed that the target and reference aDNA haplotypes are from the same population and time period. Moreover, we modelled instantaneous population growth, rather than exponential growth, experienced by most human populations. Our models were somewhat sensitive to the degree and timing of the growth. More precise modelling could suggest which aDNA targets would benefit from adding ancient haplotypes to the reference panel. However, our results suggest such detailed modelling would not change our overall conclusions.

Overall, our results caution against applying lessons from imputation of contemporary sequences to aDNA. The benefit of large reference samples is limited and low-frequency variants may not be imputed correctly. The most reliable way to improve imputation performance for ancient genomes is to add relevant ancient samples to a modern reference panel. The degree of benefit substantially depends on sample age, effective population size, and the number of available ancient genomes. Our results show that, for this reason, in the European context, European hunter-gatherers would benefit most from this approach. Future work would involve generating more high-quality ancient genomes and carefully genotyping and phasing this data. In this study, we focused on loci polymorphic in modern reference panels, as only the simulations included polymorphisms private to ancient individuals. An aDNA panel can allow access to variation that has been lost over time. In particular, such variation could provide us with a more complete view of selection acting on the human genome and also increase the imputation performance of an aDNA panel. Additionally, aDNA imputation might benefit from directly modelling the age of reference panel variants, for instance, using an ARG (Mejia-Garcia et al. 2025; Zhang et al. 2023; McClelland et al. 2026; Pope et al. 2026). This could reduce the error induced by model misspecification in aDNA imputation. It also could be beneficial to separately impute targets with the ancient and modern panels and then combine them in a meta-imputation framework (Kumar et al. 2026), to allow integration of privacy-protected modern panels and to avoid differences in genotype and phasing quality affecting reference template selection.

## Methods

### Deriving analytical imputation accuracy and solving through numerical methods

In the coalescent, mutations are independent of tree topology, and in expectation, they accrue on the tree at a rate proportional to the branch length. Therefore, we calculate the expected branch length *E[T_c_]* as a proxy for imputation error.

We obtain *E[T_c_]* under population structure and with or without adding ancient samples in a modern reference panel. Similar to Biddanda et al. (2022), we express *E[T_c_]* conditioning on the ancestral process, *A_K_(T_a_)* as in equation 1 below. *A_K_(T_a_)* represents the number of ancestral lineages remaining at *T*_a_ from a coalescent tree with size *K* at *T = 0*.

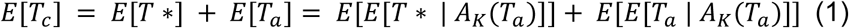

First, we consider the outer expectation, where we have *P(A_K_(T_a_) = k)*. Deriving this quantity under population structure analytically requires summing over infinitely many migration histories, which is not tractable. Therefore, we sample from the probability distribution *A_K_(T_a_)* using a continuous time markov chain (CMTC) (Sampson 2006; Wakely 2016). Furthermore, we assume that after *T_a_*, we enter the collecting phase of the coalescent, which means the structured coalescent converges to the standard Kingsman coalescent (Wakeley 2016). This assumption is standard if we assume the migration rate is constant. Therefore, we calculate the inner expectation of *E[T*]* analytically as in Biddanda et al. (2022).

We also derive *E[T_a_]*. We condition on the number of extant lineages at *T_a_*, creating a double expectation *[E[T_a_ | A_K_(T_a_)]]*. If we are using a modern reference panel, then *E[E[T_a_ | A_K_(T_a_)]] = T*_a._ If we add *d* aDNA samples to the modern reference panel, then if the aDNA target coalesces first with an aDNA reference haplotype, then *E[E[T_a_ | A_K_(T_a_)]] = 0*. Since *d + k* lines at *T_a_* are exchangeable, conditioning on *A_K_(T_a_)*, the probability of coalescing with a modern sample is 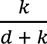. Therefore, *E[E[T_a_ | A_K_(T_a_)]]* is as follows.

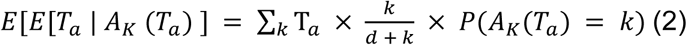

### Evaluation

We evaluate *E[T_c_]* across the following parameter settings. For reference panel sizes of n=100, 500, 1000, 3,000 and 10,000, we evaluate across deme sizes of 2, 5, and 10 with a constant migration rate of 1e-5 and 1 between demes. To test strong population structure, we evaluate the effect of 50 demes at n=3,000 with migration rates of 1e-5 and 1. We also evaluate *E[T_c_]* when adding 0,5,10, 50, 100, 500, and 1000 aDNA individuals at *T_a_* for a panel size of n=1,500, with constant migration rates 1e-5 and 1 and for migration rates as determined in a demographic model for 3 populations out of Africa: Africans (YRI), Europeans (CEU), and East Asians (CHB) (Gutenkunst et al. 2009). For this later demographic scenario, we also use the Gutenkunst et al. (2009)’s split times and generation time, and we add an instantaneous population expansion from 10,000 individuals to 10 million individuals (Si et al. 2021) at 𝜏 = 0.031 2N_e_ generations, where N_e_ is the ancestral population size of 10,000 individuals. We selected 𝜏 so that the effective population size under instantaneous growth is the same as the effective population size under exponential growth starting at time = 0.043 2N_e_ generations (Gutenkunst et al. 2009) (**Supplementary Note 1)**.

We run the CMTC 1000 times to sample from *P(A_K_(T_a_) = k).* If the panel size does not divide evenly into the number of demes, then the panel size is rounded down accordingly. For the demographic model, there are stopping times for each split time and the instantaneous population growth. We run the CMTC once to obtain the final state after each stopping time and then, for the remainder of the time after all stopping times, run the CMTC 1000 times to sample from *P(A_K_(T_a_) = k)*.

### Model misspecification for imputing aDNA with modern reference panels

Model misspecification of the Li and Stephens model occurs when there is more than one best template in the reference panel. We derive the number of best templates, *u*, when imputing an aDNA target with a modern reference panel. We condition on the number of lineages, *m_c_* and *m_a_*, extant on the tree at *T_c_* and *T_a_*, respectively.

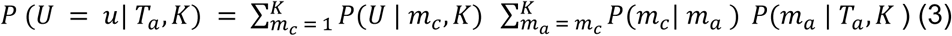

Here, 𝑃(𝑚_𝑐_| 𝑚_𝑎_) = 𝑃(𝐼_𝑗_) for *j = m_c_ +1*, where 𝑃(𝐼_𝑗_) represents the probability that the aDNA target coalesces at the *j → j-1* coalescent event (Biddanda et al. 2022). We derive 𝑃(𝑈 = 𝑢 | 𝑚_𝑐_, 𝐾) by treating the problem as one of integer composition (**Supplementary Note 1**). We then calculate 𝑃 (𝑈 = 𝑢| 𝑇_𝑎_, 𝐾) for *K*=6000 and *K*=200,000 at *T_a_* ∈ {0.004, 0.04, 0.02, 0.1} and additionally at *T_a_* ∈ {0.0002, 0.0004, 0.00002, 0.00004, 0.000002, 0.000004} for *K*=6000. We multiply *T_a_* by 2N_e_ generations * 25 years per generation to obtain time in years before present.

### Coalescent-based simulations

#### msprime simulations

We used msprime (Baumdicker et al. 2022) v1.3.4 to produce genotypes on chromosome 1 for sampled individuals from a simulated genealogy. We used the HapMap recombination map (International HapMap 3 Consortium et al. 2010) and a mutation rate of 1.25e-8 per generation and base pair. The model underlying the simulations is a modified version of the ‘multi-population model of ancient Eurasia’ (‘AncientEurasia_9K19’) available through the stdpopsim python package and based on (Kamm et al. 2019). We removed populations not directly related to the populations that we simulated. The demographic events and effective population sizes common to all simulations are in **Supplementary Table 2** and **Supplementary Table 3**, respectively.

We sampled individuals from three populations at t=0: Mbuti-like, Han-like, and Sardinian-like. The imputation reference panels we used had a fixed number of Han-like and Mbuti-like genomes (500 from each population) and a varying number of ‘Sardinians’ between 100 and 10,000.

For the first set of simulations, all ancient individuals come from the ‘Sardinian/LBK’ branch with a fixed N_e_ of 15,000. In order to eliminate additional demographic processes, we removed the ‘WHG’ admixture pulse into the ‘Sardinians’ and modified the LBK N_e_ to match the Sardinians’ before the split of these two populations. We sampled 500 individuals at t=165 generations, Ref165, and at t=750 generations, Ref750, to serve as reference (**Supplementary Fig. 5**). To be noted, the imputation reference panels only had ancient haplotypes belonging to either of these two datasets at a time. We sampled ten ancient targets at five different time points: 40, 80, 250, 500 and 1000 generations in the past (**Supplementary Table 4**).

In the second set of simulations, we created two populations splitting from ‘LBK’ and ‘WHG’, from which we sampled the ancient individual samples to be the imputation targets: ‘LBK0’ and ‘WHG0’ splitting from ‘LBK’ and ‘WHG’ at t=600 generations, respectively, and ‘LBK1’ and ‘WHG1’, splitting from ‘LBK’ and ‘WHG’ at t=420 generations, respectively. The ancient DNA panels containing 500 individuals each are sampled directly from ‘LBK’ and ‘WHG’ (**Supplementary Table 5**). We additionally varied the effective population size of ‘LBK’ before the split that led to the Sardinian-like branch: 1) constant population size of 12,000 (Kamm et al. 2019), 2) constant population size of 100,000, and 3) instant expansion from 12,000 to 100,000 at t=430 generations before present.

#### Reads simulations

We used NGSNGS (Henriksen et al. 2023) to simulate reads for the target ancient genomes with parameters *-seq SE -ld norm,350,20 -q1 NGSNGS/Test_Examples/AccFreqL150R1.txt -cl 100*. The required reference genome in fasta format was generated using as template the hs37d5 assembly and by introducing the reference allele from the simulated variants. This step produced bam files with genome coverage of 1x.

#### Population structure

To examine the population structure of the simulated genomes, we calculated Fst (fixation index) (Wright 1951) using ADMIXTOOLS2 (Maier et al. 2023) and performed MDS (multidimensional scaling). For the MDS, we limited the number of ‘Sardinian’ genomes to 500 (from a total of 10,000), as MDS is sensitive to population sample size. We generated a distance matrix whose metric was 1-IBS (identity by state) using plink (Chang et al. 2015) v1.90b7.7_20241022 and the parameters -*-maf 0.01 --distance square 1-ibs flat-missing*. The MDS coordinates were then calculated with the R (v4.5.2) function cmdscale with 10 components having as input the distance matrix.

In some simulations, the two first MDS coordinates indicate some substructure for some populations, and there is little distinction between the ‘WHG’ target and reference genomes and between LBK-related individuals and the ‘Sardinians’. Nonetheless, the Fst estimates confirm that we were able to simulate the intended genealogies across the different simulation sets (**Supplementary Fig. 6** and **Supplementary Fig. 7**).

#### Imputation and imputation assessment

We imputed the low-coverage genomes using GLIMPSE2 (Rubinacci et al. 2023) and default parameters. We first generated chunks for the simulated reference panels using GLIMPSE2_chunk and then split the reference panel into those chunks with GLIMPSE2_split_reference. For each chunk, we imputed with GLIMPSE2_phase and subsequently ligated the imputed chunks into a full chromosome with GLIMPSE2_ligate. To assess imputation performance, we used GLIMPSE2_concordance.

In our analysis, we mainly use two metrics to assess imputation performance: 1) non-reference discordance (NRD), and 2) squared Pearson correlation, *r^2^*, between imputed dosages and ground truth genotypes. NRD is the ratio of the total number of errors and the total number of considered sites minus the correctly imputed homozygous reference allele sites.

#### Real ancient DNA imputation panel

To investigate how including ancient haplotypes in a reference panel affects imputation in reality, we gathered 95 high-quality ancient genomes, with depth of coverage>11.5x (median of 26.5x, **Supplementary Table 7**), from the past 45,000 years. We explored how these genomes relate to present-day populations via PCA (principal component analysis, **Supplementary Fig. 17**). We used smartpca (Patterson et al. 2006) to project (*lsqproject: YES*) the ancient data into the two first PCs defined by 1) the 1000 Genomes panel data 2) a subset of Western Eurasian individuals from the Human Origins dataset.

#### Read simulation with aDNA-like damage

To test different genotype calling approaches (Supplementary Note 4), we used gargammel (Renaud et al. 2017) to simulate ancient-like reads. We used as template the genome of individual NA20502 from 1000 Genomes and generated deaminated reads with 20x average coverage for chromosome 1 without contamination. We followed (Koptekin et al. 2025) to specify deamination parameters. We ran the following command: *gargammel.pl -c 20 --comp 0,0,1 --loc 4.106487474 --scale 0.358874723 --minsize 30 --maxsize 150 -damage 0.024,0.36,0.0097,0.55*. We then removed adapters with AdapterRemoval v2.3.4 (Schubert et al. 2016), and mapped the reads with bwa v0.7.18 (Li and Durbin 2009), following aDNA mapping best practices (*bwa aln -l 1024 -n 0.01 -o 2*).

#### Genotype calling high coverage ancient genomes

Before genotype calling, we applied bamRefine (Koptekin et al. 2025) to mask 1000 Genomes polymorphisms that are prone to be affected by postmortem damage within a set distance from the ends of the reads. This distance was determined based on the C-to-T (5’ end) and G-to-A (3’ end if libraries were double stranded) substitution rates (postmortem damage), that we estimated with PMDtools (Skoglund et al. 2014). We then called genotypes with bcftools at the 1000 Genomes biallelic SNPs for chromosome 1 and required a minimum base quality of 20 and mapping quality of 30. Finally, we removed depth outliers (minimum depth of 10 and maximum equal to twice the average chromosome coverage) and variants with QUAL<30 (see **Supplementary Note 4** for an assessment of genotype calling approaches for high-quality ancient genomes).

#### Assembling the aDNA high-quality dataset

We found that imputation performance improved when the high coverage ancient calls, i.e., in the reference panel, were also imputed (**Supplementary Fig. 21**), compared to directly using the bcftools genotype calls. Therefore, we imputed genotype likelihoods for the high coverage ancient genomes using GLIMPSE2 with default parameters and the 1000 Genomes as a reference panel. Afterwards, we phased the imputed ancient calls with SHAPEIT5 (Hofmeister et al. 2023) using again the 1000 Genomes as a reference panel (for an assessment of phasing switch error rates in ancient genomes, see **Supplementary Note 5**). Finally, we merged the phased ancient dataset with 1000 Genomes.

To test imputation performance, we applied a leave-one-out approach. For each ancient genome in the dataset, we removed it from the reference panel and we downsampled the raw reads without masking with bamRefine, to 1x average coverage. We then imputed this downsampled genome with GLIMPSE2 as before, but using a reference panel consisting of 1000 Genomes and n-1 ancient genomes.

#### Relate variant age

We used the SNPs ages as estimated in (Speidel et al. 2019) for the CEU (Utah residents with Northern and Western European ancestry) individuals in 1000 Genomes using Relate. We split the variants into age bins (age in generations before present): 0-200, 200-400, 400-800, ≥800. For a variant to belong to a specific bin, its midpoint age must be within the bounds of that bin and its lower bound age must be greater or equal than half the lower bin limit.

### Datasets

#### High coverage 1000 Genomes

We used a GRCh37/hg19 version of the 30x 1000 Genomes phase 3 (Byrska-Bishop et al. 2022), that was lifted over in the context of the study (Akbari et al. 2026).

#### Human Origins

We used 1070 individual samples from the Human Origins (Western Eurasians) dataset to perform PCA (Broushaki et al. 2016; Patterson et al. 2012; Lazaridis et al. 2014, 2016; Reitsema et al. 2022).

#### High coverage ancient genomes

Our aDNA reference panel contains 95 publicly available high-quality genomes (**Supplementary Table 7**) (Wohns et al. 2022; de Barros Damgaard et al. 2018; Rohland et al. 2022; Sato et al. 2021; Simões et al. 2024; Valdiosera et al. 2018; Childebayeva et al. 2024; Immel et al. 2021; Simões et al. 2023; Lipson et al. 2022; Fernandes et al. 2021; Sikora et al. 2019; Flegontov et al. 2019; Gokhman et al. 2020; Gamba et al. 2014; Moreno-Mayar et al. 2018; Fu et al. 2014; Lazaridis et al. 2014; Rasmussen et al. 2014, 2010; Rodríguez-Varela et al. 2023; Marchi et al. 2022; Akbari et al. 2026).

## Acknowledgements

We thank Robin J. Hofmeister, Romain Fournier, Alison Barton, Vincent de Bakker, Ali Akbari, Gideon Bradburd and the Bradburd and Zöllner labs for discussions. S.Z. and K.H.K. were supported by NIH grant R01HG005855 to S.Z. B.S.d.M. was supported by an EMBO Postdoctoral Fellowship (ALTF 53-2025). B.S.d.M. and D.R. were supported by NIH grant HG012287. D.R.’s work was supported by the Allen Family Philanthropies; by John Templeton Foundation grant 61220; and by the Howard Hughes Medical Institute (HHMI).

## Author contributions

B.S.d.M., K.H.K., D.R., and S.Z. developed the ideas. B.S.d.M. and K.H.K. completed the analyses and wrote the manuscript with input from D.R. and S.Z..

