## Supplementary Information for "Optimal Reference Panel Design in Ancient DNA Imputation from Coalescent Theory, Simulation, and Real Data Application with an Ancient Reference Panel"

### Supplementary Information File

**Supplementary Note 1 – Analytical Modeling** **1**
**Supplementary Note 2 – Coalescent-based simulations** **7**
**Supplementary Note 3 – An aDNA reference panel** **20**
**Supplementary Note 4 – Testing genotype calling approaches for aDNA** **21**
**Supplementary Note 5 – Assessing phasing switch error rate** **26**
**Supplementary Note 6 – Variant age and allele frequency effect on imputation** **29**
References 30

### Supplementary Note 1 – Analytical Modeling

*Model misspecification*

We derive $P(U = u \mid m_c, K)$ by treating this problem as having *K* objects to be composed into $m_c$ pieces, such that each piece must have a positive number of objects. WLOG, we assume that the aDNA target coalesces with the leftmost branch of the tree. There are $\frac{K-1}{m_c - 1}$ total possible compositions for the leftmost piece, which serves as our denominator. For the numerator, we place *u* objects in the leftmost piece. Now, we have *K- u - 1* objects that must be divided into $m_c$ *- 1* pieces, which yields $\frac{K-u-1}{m_c - 2}$ possible compositions.

$$P(U = u|m_c, K) \begin{cases} \dfrac{\frac{K - u - 1}{m_c - 2}}{\frac{K - 1}{m_c - 1}}, & if \quad 2 \le m_c \le K - u + 1 \\ 1 & if \; mc = 1 \; and \; U = K \\ 0 & \text{if} \; mc > K - U + 1 \end{cases}$$

*Probability of the number of best templates*

The probability distribution of the number of best templates depends on $T_a$ and the reference panel size. As the number of best templates increases, the probability decreases rapidly for $T_a$ = 40 generations. However, for $T_a$ = 400 or 1000 generations the probability decreases much less sharply until the number of best templates equals the number of haplotypes in the reference panel. This represents the situation where all modern reference panel haplotypes reach their most recent common ancestor more recently than $T_a$. The shape of this probability distribution is roughly the same regardless of panel size; however, the axes' scale differs substantially (**Supplementary Fig. 3**).

*Calibrating instantaneous population growth*

We select a time $\tau$ for instantaneous population growth from $N_0$ to $N_1$ such that harmonic mean of the effective population sizes is equal to the harmonic mean of effective population sizes under exponential growth as parametrized in Gutenkunst et al. (2009). We assume $N_0$ = 10,000 individuals and $N_1$ = 10 million individuals (Si et al. 2021).

We solve the following equation, where the left-hand size represents instantaneous population growth and the right hand side represents exponential population growth with $b$ = 0.004/ generation as defined for CEU (Utah residents with Northern and Western European ancestry in 1000 Genomes) (Gutenkunst et al. 2009), which we convert to coalescent units by multiplying by $2N_0$. Here, $t_g$ represents the start of growth in the Gutenkunst et al. (2009) model, which is at t=0.0434 in units of $2N_0$ generations.

$$\frac{t_g}{(1/2N_1)\times\tau + (1/2N_0)(t_g - \tau)} = \frac{t_g}{\int_0^{t_g} 1/(2N_0\, e^{\,bt})dt}$$

The solution of this equation is $\tau$ = 0.0303509 in units of $2N_0$ generations.

We also calibrate instantaneous population growth by fixing $\tau = t_g$ and solving for $N_1$, using the above equation. We obtain $N_1$ = 34,753 individuals from this approach and plot our results in **Supplementary Fig. 4**

*Theoretical model assumptions*

For our theoretical model, we make several assumptions that are standard in coalescent theory. First, we assume that the sample size is much smaller than the effective population size, which may be violated in our examination of large reference panel sizes because the effective population size is not necessarily much larger than the sample size. However, this means that multiple lineages are more likely to coalesce in one event, yielding a smaller number of lineages extant at $T_c$ than in our model (Bhaskar et al. 2014). Thus, accounting for these likely violations of our model will only strengthen our conclusions that aDNA targets do not benefit much from larger reference panels, and, in fact, may benefit more from an ancient reference panel. Furthermore, we assume that the coalescent rate is much smaller than the migration rate at the time of the ancient sample, which is not the case for young aDNA targets. However, our results and recommendations are more focused on older samples, for which this assumption is reasonable. Finally, in the theoretical work we consider a single locus which is standard in such models (Jewett et al 2013; Si et al 2021); moreover, the simulated results account for recombination.

**Supplementary Table 1:** Probabilities of the Li and Stephens model being correctly specified (i.e. having one best template) across aDNA target age (years before present) assuming Ne=10,000 individuals and generation time of 25 years) and reference panel sizes of n=3,000 and n=100,000.

| Probability | Number of Best Templates (u) | Ancient Sample Age (Ta) Years Before Present | Reference Panel Size (n) |
|---|---|---|---|
| 0.0512 | 1 | 2,000 | 3,000 |
| 0.0109 | 1 | 10,000 | 3,000 |
| 0.0055 | 1 | 20,000 | 3,000 |
| 0.0022 | 1 | 50,000 | 3,000 |
| 0.0017 | 1 | 2,000 | 100,000 |
| 0.0003 | 1 | 10,000 | 100,000 |
| 0.0002 | 1 | 20,000 | 100,000 |
| 0.0001 | 1 | 50,000 | 100,000 |

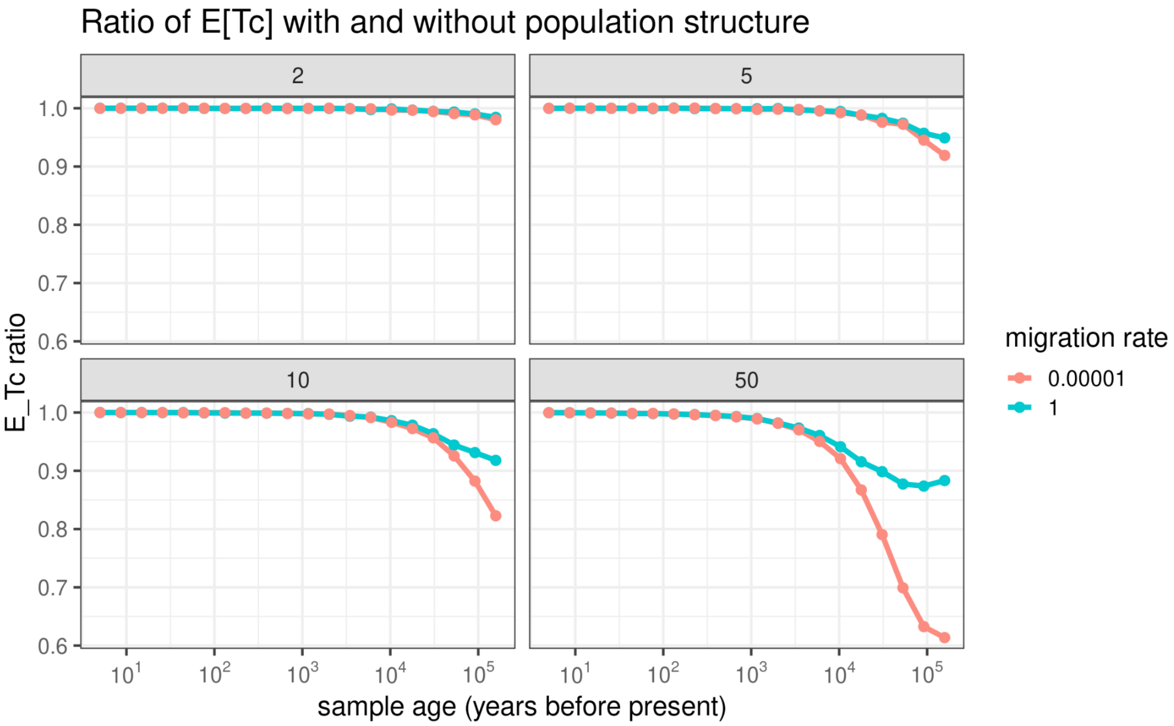


**Supplementary Fig. 1:** Ratio of the expected coalescent times between aDNA target and modern templates in structured versus unstructured population against target age in years (assuming Ne=10,000 individuals and a generation time of 25 years) for different deme sizes and migration rates with a panel size of 3000 individuals. Each panel represents a different number of demes (2, 5, 10, 50), and the color of the line within a panel represents a migration rate. A migration rate of 1 is “high” whereas a migration rate of 1e-5 is “low.”

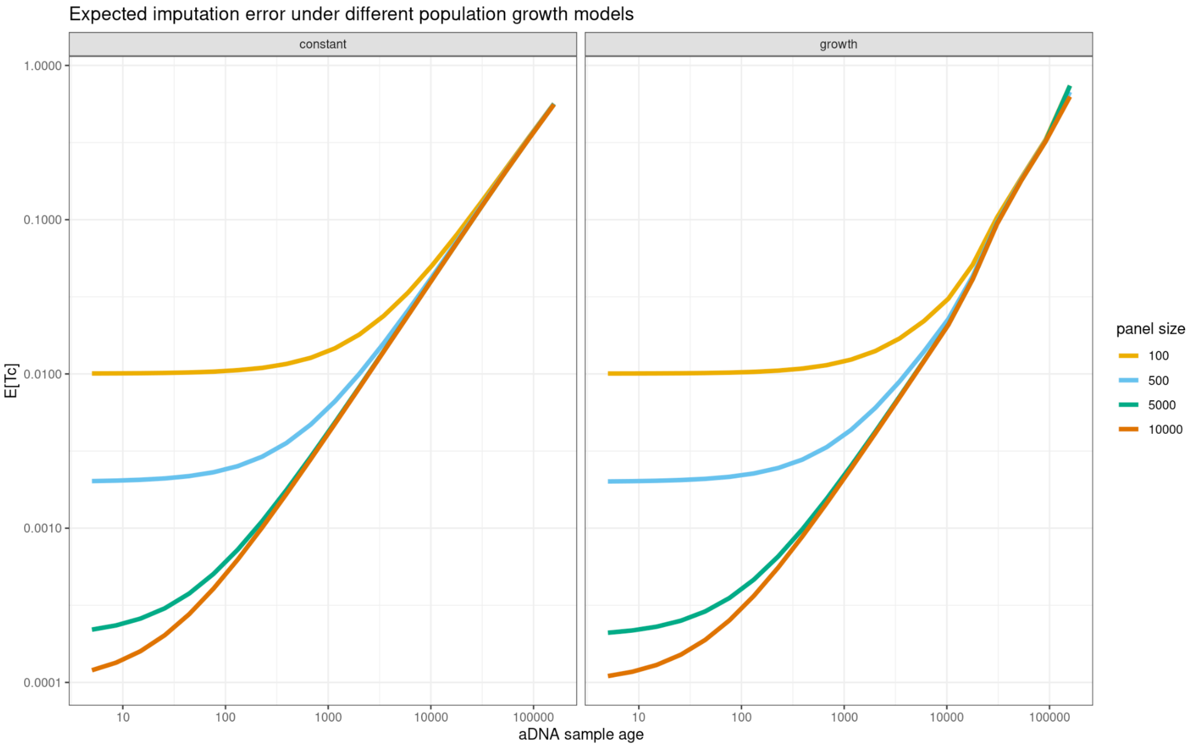


**Supplementary Fig 2:** For the instantaneous growth model, we plot the age of the ancient sample in years before present against the expected coalescence time between templates and target (i.e. E[Tc]) for four different reference panel sample sizes (n=100, 500, 5000, 10000). We compare it to the constant model here in the left panel, which is also displayed in the main text as Figure 2a.

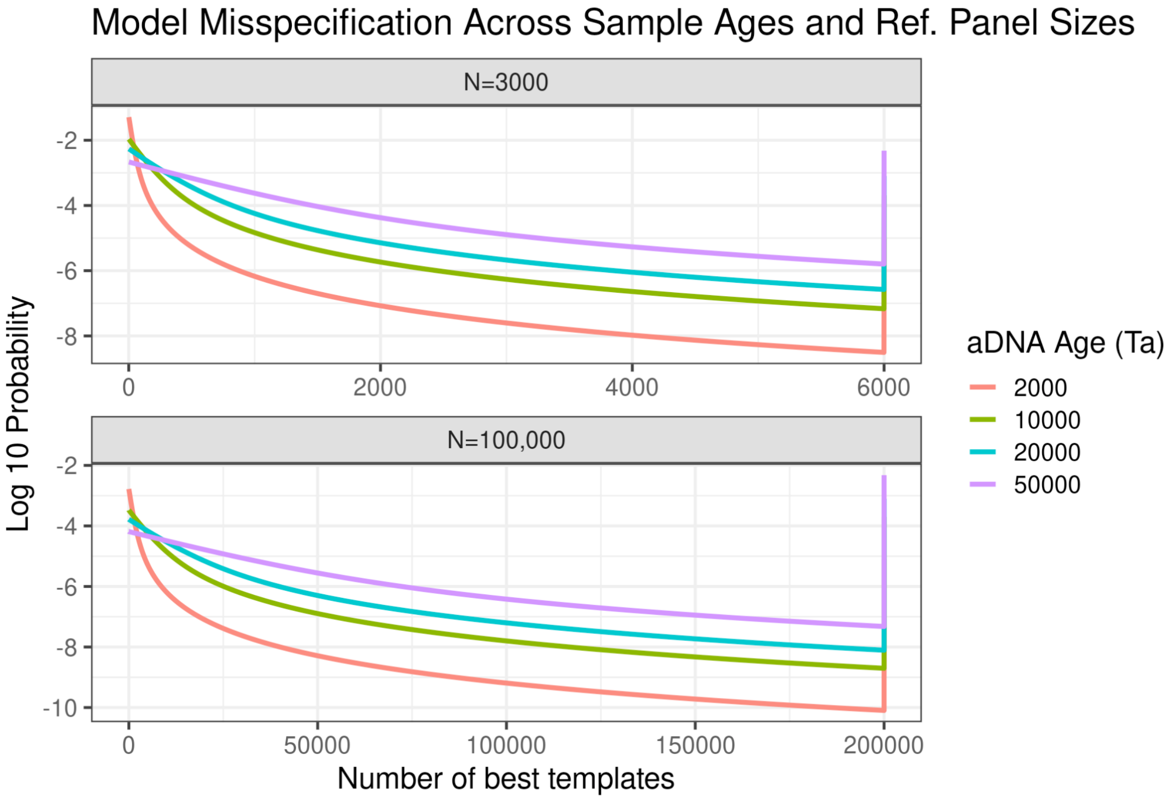

**Supplementary Fig.3:** Probability distribution of best templates across four ancient sample times (2,000, 10,000, 20,000 and 50,000 years before present, assuming Ne=10,000 individuals and generation time of 25 years) and two reference panel sizes (n=3000, n=100,000), representing the approximate sizes of the 1000 Genomes panel and the TopMed panel, respectively.

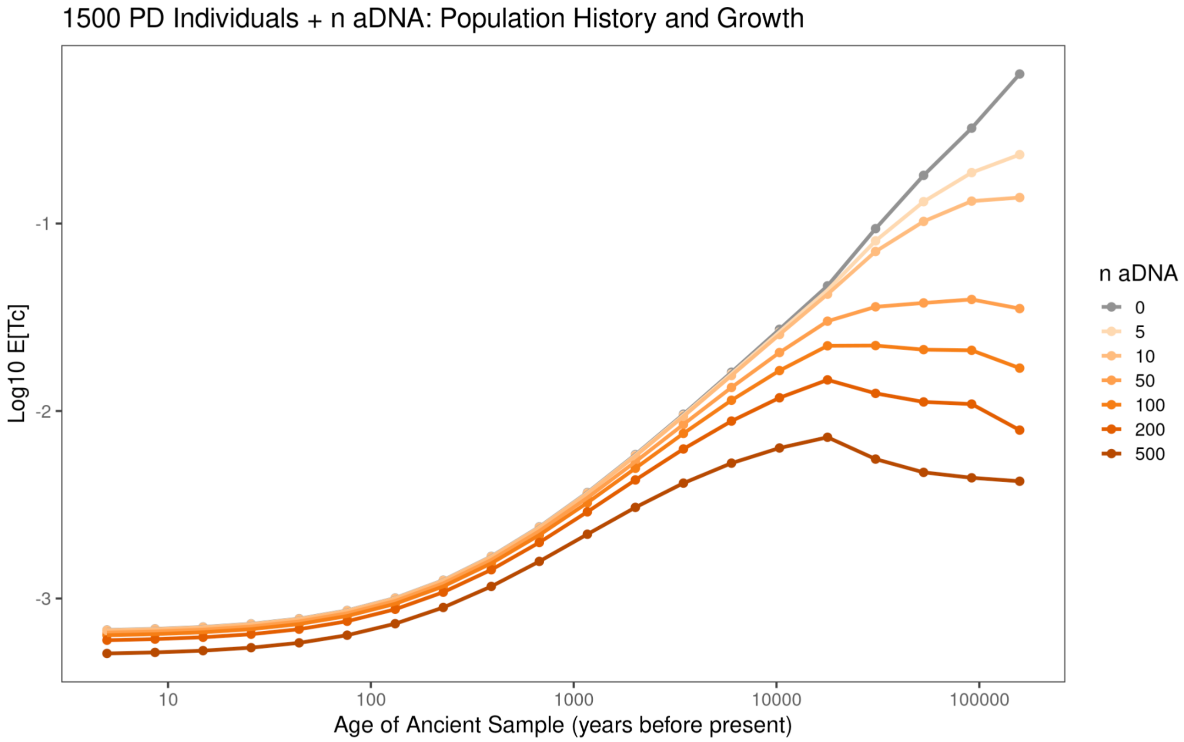


**Supplementary Fig. 4:** The expected coalescence time between target and best templates for aDNA imputation with 1,500 present day (PD) individuals and between 0 to 500 ancient samples (n aDNA) in a reference panel across aDNA target times (years before present). Here, we have instantaneous population growth from Ne=10,000 individuals to Ne=34,753 individuals at the time when growth starts for Europeans in Gutenskunst et al. (2009).

### Supplementary Note 2 – Coalescent-based simulations

#### Simulations parameters

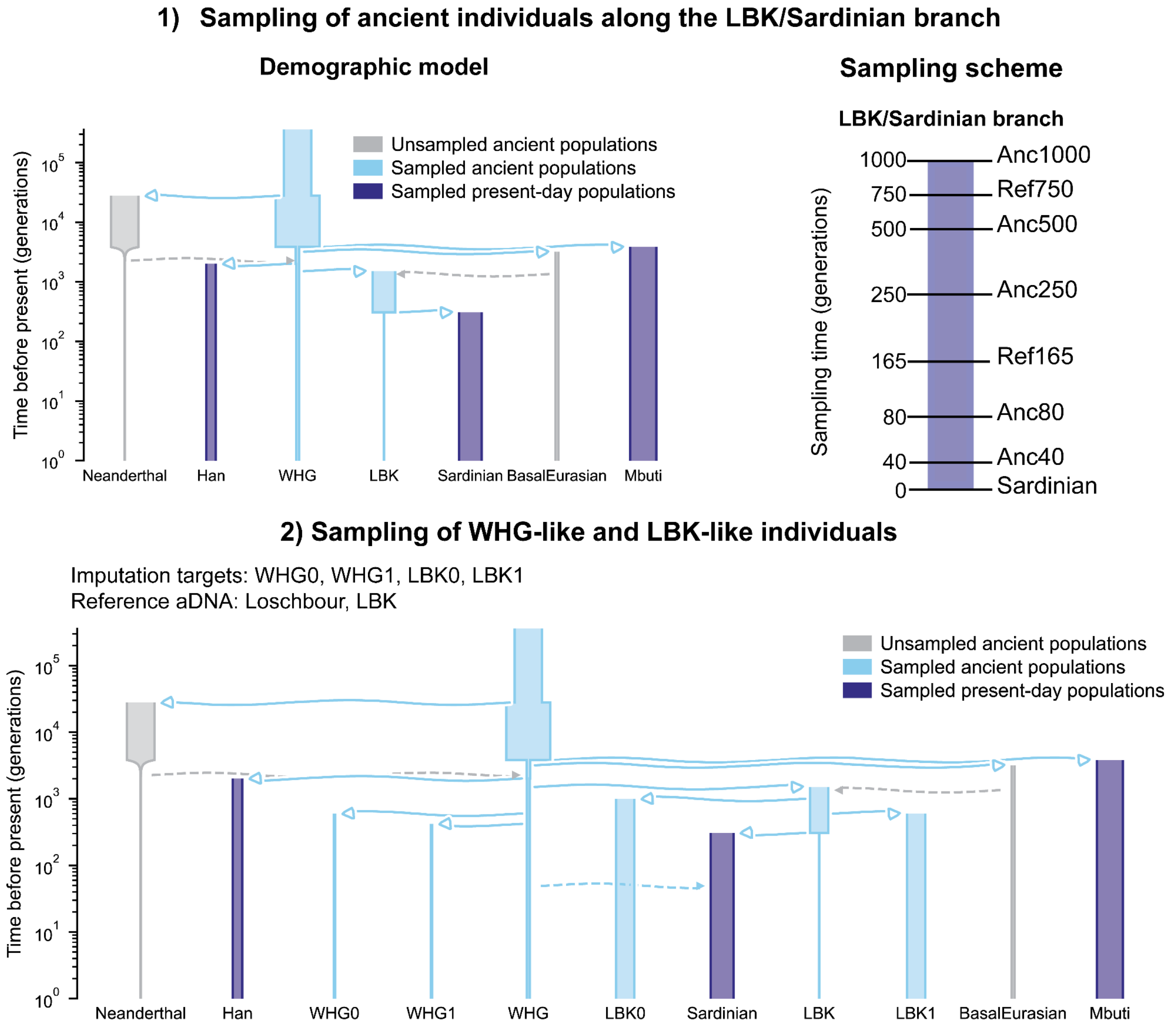


**Supplementary Fig. 5:** Demographic models underlying the coalescent-based simulations. Top: on the left, we show the demographic model underlying simulations where we sampled ancient chromosomes from the 'LBK/Sardinian' branch, whose sampling time is illustrated on the right side. 'RefX' stands for reference panel sampled at t=X generations in the past and 'AncY' stands for target genomes to be imputed and sampled at t=Y generations before the present. Bottom: demographic model that we used to simulate ancient individuals from LBK-like and WHG/Loschbour-like populations. Both models were based on (Kamm et al. 2019), but we modified the model on top by removing an admixture pulse from 'Loschbour' into the 'Sardinian' population and by changing the effective population of 'LBK' from 12,000 to 15,000 before the split into 'Sardinian'. The solid arrows represent population splits and the dashed arrows represent admixture pulses.

**Supplementary Table 2:** Demographic events and their time of occurrence in the model underlying the set of simulations from which we sampled WHG and LBK individuals. The events and parameters follow (Kamm et al. 2019).

| Event | Time (generations before present) |
|---|---|
| Split between modern humans and Neanderthals | 27840 |
| Split between African and non-Africans | 3832 |
| Split between Basal Eurasians and other non-Africans | 3192 |
| Admixture pulse from Neanderthals into non-Africans (2.96%) | 2272 |
| Split between Han and other non-Basal Eurasians | 2016 |
| Split between LBK and other non-Basal Eurasians | 1508 |
| Admixture pulse from Basal Eurasians into LBK (9.36%) | 1348 |
| Split between LBK and Sardinians | 307.6 |
| Admixture pulse from Loschbour (WHG) into Sardinians (3.17%) | 49.2 |

**Supplementary Table 3:** Effective population sizes (Ne) (Kamm et al. 2019) of modelled populations. These parameters remained unchanged across simulation sets and versions.

| Population | Effective population size (Ne) (time in generations) |
|---|---|
| WHG | 0≤t<2016: 1,920<br>2016≤t<3832: 2,340<br>3832≤t<27840: 29,100<br>t≥27840: 18,200 |
| Neanderthal | 0≤t<2000: 86.9<br>2000≤t<3,832: 18,200.000000000007<br>t≥3832: 18,200 |
| Mbuti | 17,300 |
| Basal Eurasian | 1,920 |
| Han | 6,300 |

**Supplementary Table 4:** Effective population size (Ne), sampling time and sample size for the first set of simulations, where we sampled individuals from the Sardinian/LBK lineage at different points in the past. For the sampled ancient individuals, we indicate between brackets the source population in simulations. We modified some of the parameters of (Kamm et al. 2019): we removed the admixture pulse from Loschbour (WHG) into Sardinians and we set the initial Ne for LBK equal to the Sardinian Ne before t=307.6 generations (split time between LBK and Sardinians).

| Population | Effective population size (Ne) | Sampling time (generations before present) | Sample size |
|---|---|---|---|
| Mbuti | 17,300 | 0 | 500 |
| Han | 6,300 | 0 | 500 |
| Sardinian | 15,000 | 0 | 10,000 |
| Ref165 (Sardinian) | 15,000 | 165 | 500 |
| Ref750 (LBK) | 15,000 | 750 | 500 |
| Anc40 (Sardinian) | 15,000 | 40 | 10 |
| Anc80 (Sardinian) | 15,000 | 80 | 10 |
| Anc250 (Sardinian) | 15,000 | 250 | 10 |
| Anc500 (LBK) | 15,000 | 500 | 10 |
| Anc1000 (LBK) | 15,000 | 1000 | 10 |

**Supplementary Table 5:** Effective population size (Ne), sampling time and sample size for the second set of simulations, where we sampled individuals from ‘LBK’ and ‘Loschbour/WHG’ deeper in time. The effective population sizes shown below correspond to the simulations whose results we show in the main text. We have two more versions of this set of simulations where we modified Ne of ‘Sardinians’ and/or ‘LBK’.

| Population | Effective population size (Ne) | Sampling time (generations before present) | Sample size |
|---|---|---|---|
| Mbuti | 17,300 | 0 | 500 |
| Han | 6,300 | 0 | 500 |
| Sardinian | 15,000 | 0 | 10,000 |
| WHG | 1,920 | 500 | 500 |
| WHG0 | 1,920 | 500 | 5 |
| WHG1 | 1,920 | 320 | 5 |
| LBK | 12,000 | 500 | 500 |
| LBK0 | 12,000 | 500 | 5 |
| LBK1 | 12,000 | 320 | 5 |

#### Population structure and characteristics

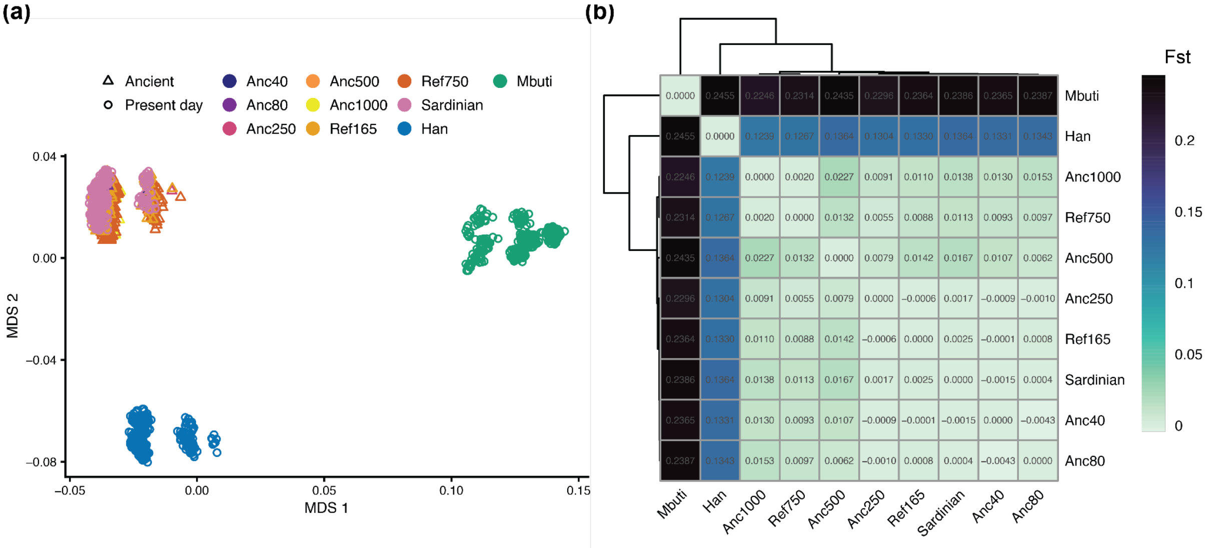


**Supplementary Fig. 6** Population structure for the simulations with sampling across time on the Sardinian/LBK branch: a) coordinates for the two first MDS for 500 “Sardinians” and all individuals from the other populations; b) heatmap of Fst estimates across all simulated populations.

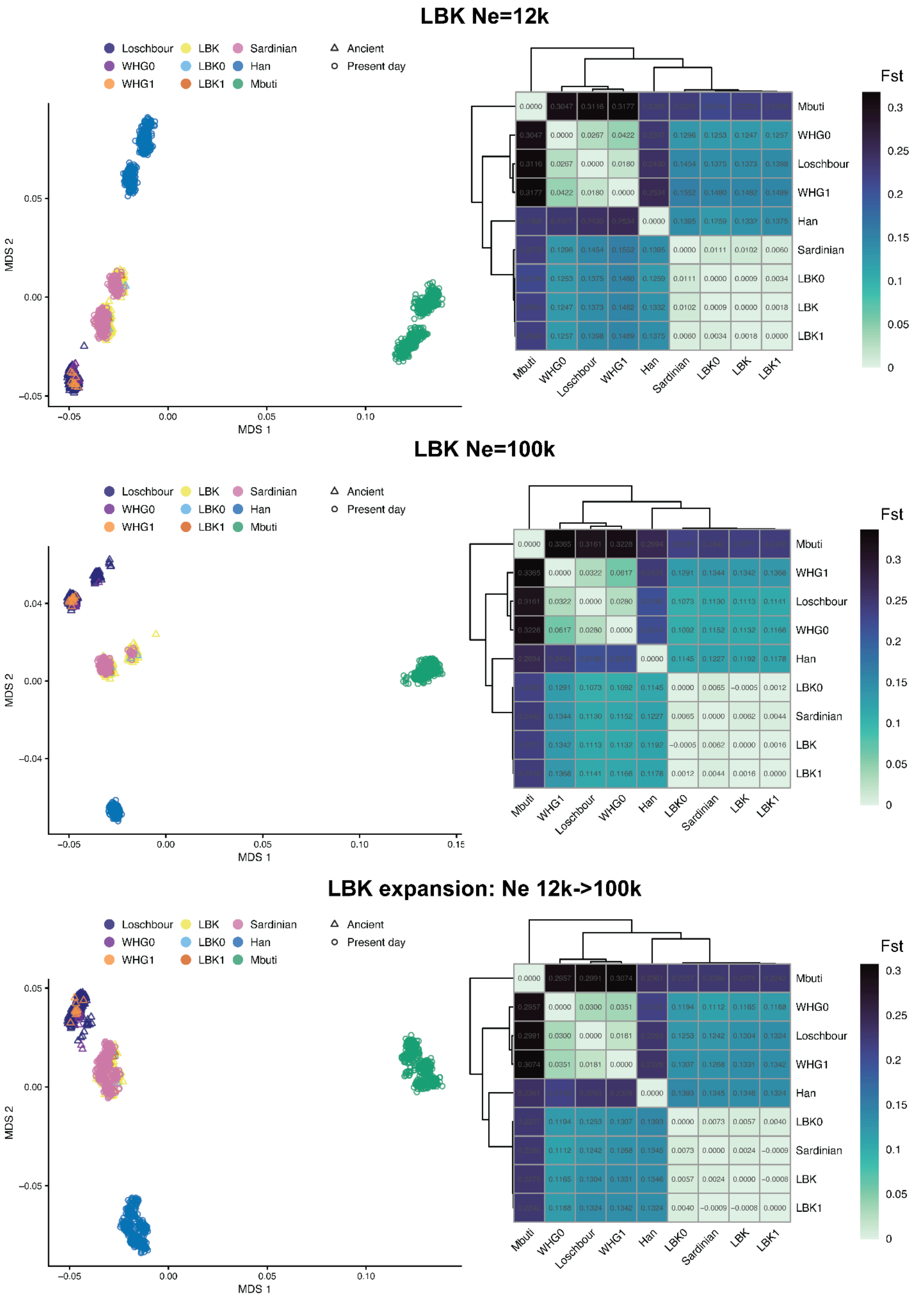


**Supplementary Fig. 7:** Population structure for the simulations with sampling of contemporaneous ‘LBK’ and ‘WHG’ individuals. We show population structure for three versions of the simulations, where we changed effective population size (Ne) for the LBK-like populations, from top to bottom: Ne=12,000, Ne=100,000, and Ne increases from 12,000 to 100,000 at t=430. Coordinates for the two first MDS are shown on the left and tables with Fst estimates for all populations are depicted on the right.

#### Allele frequency distribution

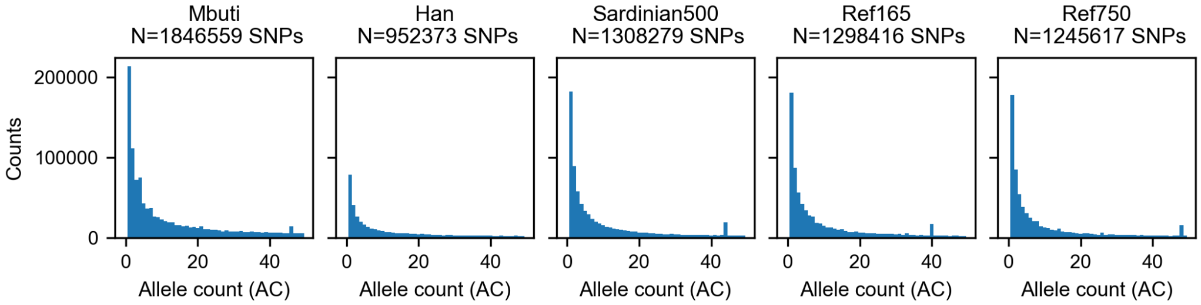


**Supplementary Fig. 8:** Allele counts per reference population (500 individuals each) in time varying sampling simulations. On top of each subplot, we show the number of bi-allelic SNPs (AC>0 and AC<1000) per population.

**Supplementary Table 6:** Number of variants in 500 present-day 'Sardinians' ('Sardinian500'), 500 ancient individuals from the 'Sardinian' lineage 165 generations before present (t=165, 'Ref165'), and 500 ancient individuals from the Sardinian lineage 750 generations before present (t=750, 'Ref750'), and how they are shared across the different groups.

| | **Ref750** | **Ref165** | **Sardinian500** |
|---|---:|---:|---:|
| **Ref750** | 399,633 | 843,617 | 813,575 |
| **Ref165** | 843,617 | 246,227 | 1,019,780 |
| **Sardinian500** | 813,575 | 1,019,780 | 286,132 |

#### Imputation results

##### Simulations with sampling from the same lineage at different time points

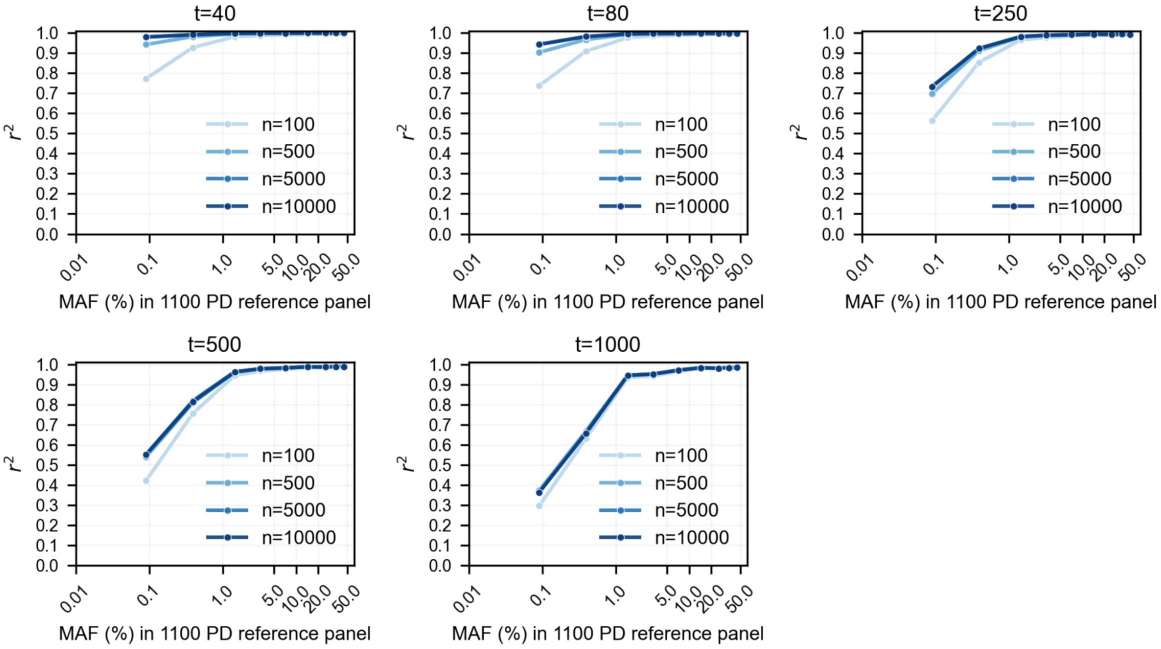


**Supplementary Fig. 9:** Effect of varying the number of present-day Sardinians in the reference panel on imputation of ancient genomes with varying ages. The different plots depict aggregate Pearson squared correlation, $r^2$, between imputed genotype dosages and ground truth genotypes as a function of minor allele frequency, MAF, as estimated from the reference panel containing 1,100 present-day individuals ('1100 PD'). Each plot represents imputation accuracy for a sample size of 10 individuals with the same ancestry and age, which increases across the plots from left to right and top to bottom.

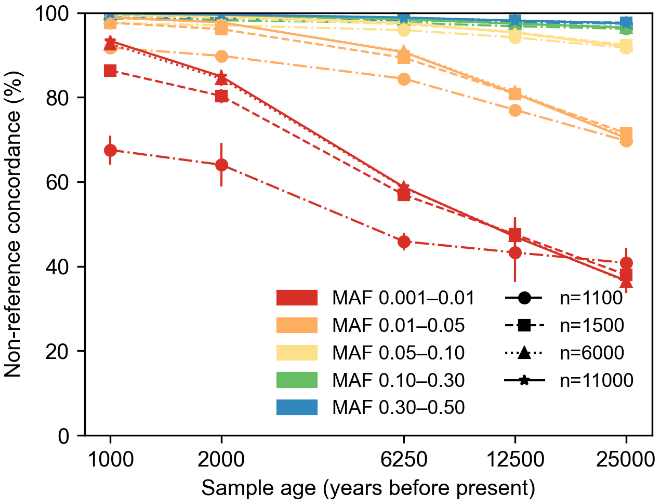


**Supplementary Fig. 10:** Effect of reference panel size on imputation of ancient genomes stratified by sample age and minor allele frequency (MAF). We plot mean non-reference concordance (NRC) across samples of the same age as a function of sample age in years before present (assuming 1 generation = 25 years). NRC was calculated as 1-NRD (non-reference discordance). Colors indicate different MAF bins, lines and markers correspond to different panel sizes (varying between 1,100 and 11,000), and error bars indicate twice the standard error of NRC. MAF was estimated in the largest reference panel containing 11,000 individuals (10,000 ‘Sardinians’).

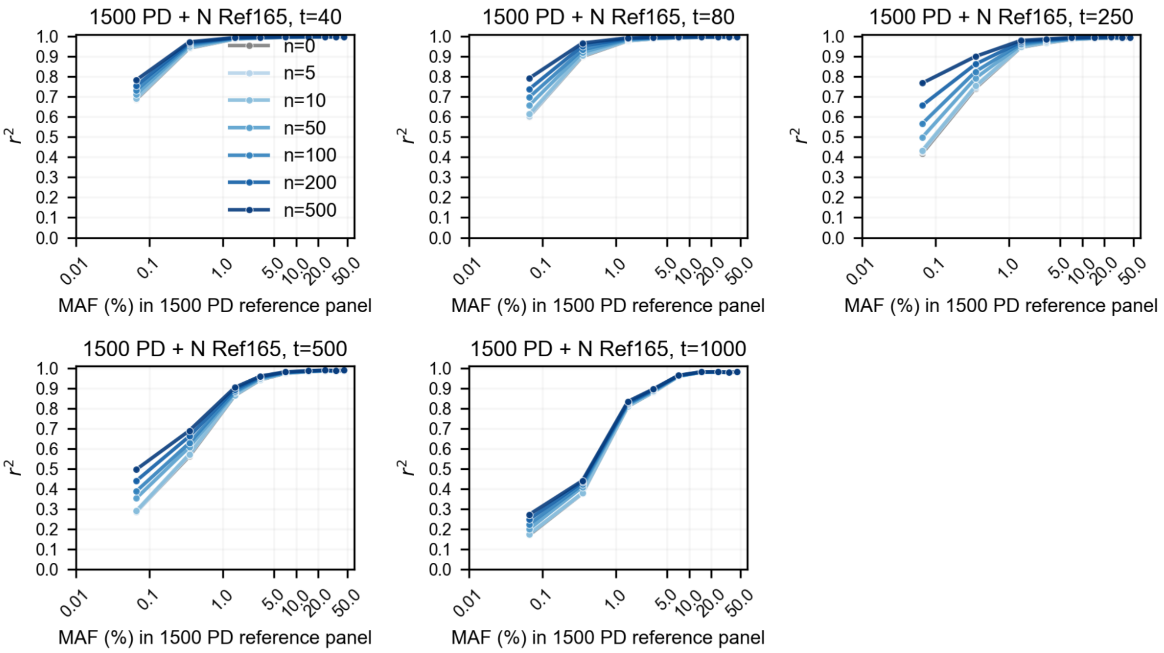


**Supplementary Fig. 11:** Effect of varying the number of ancient haplotypes sampled at 165 generations in the past (‘Ref165’) in the reference panel on imputation of ancient genomes with varying ages. The different plots depict aggregate Pearson squared correlation, $r^2$, between imputed genotype dosages and ground truth genotypes as a function of minor allele frequency, MAF, as estimated from the reference panel containing 1,100 present-day individuals (‘1100 PD’). Each plot represents imputation accuracy for a sample size of 10 individuals with the same ancestry and age, which increases across the plots from left to right and top to bottom.

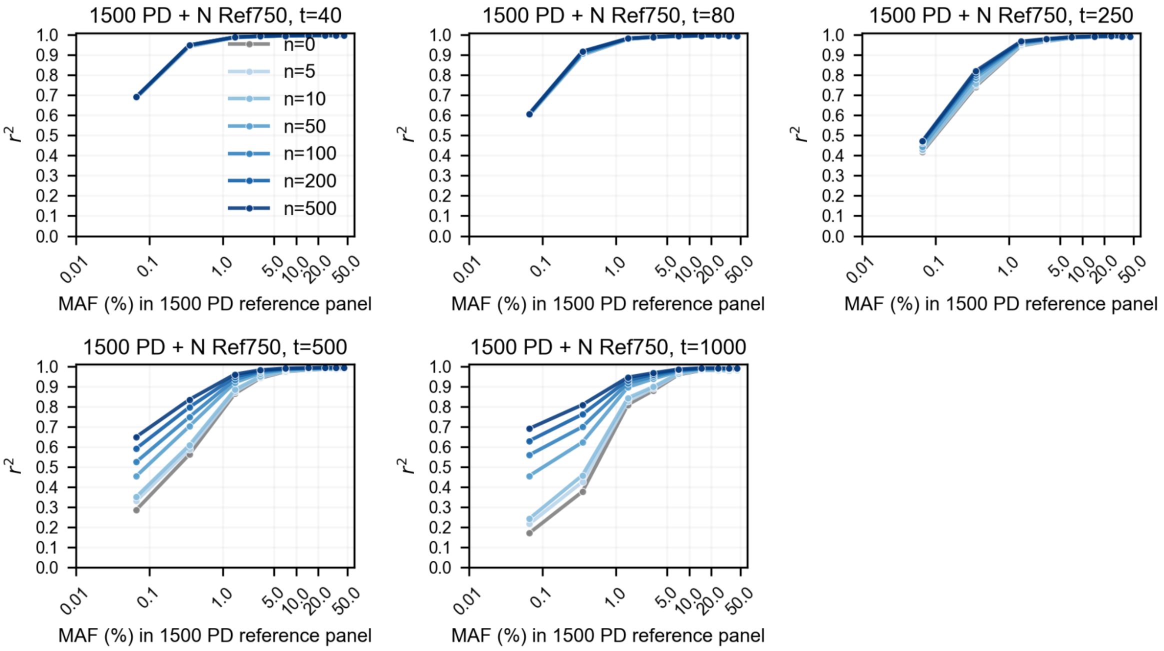


**Supplementary Fig. 12:** Effect of varying the number of ancient haplotypes sampled at 750 generations in the past ('Ref750') in the reference panel on imputation of ancient genomes with varying ages. The different plots depict aggregate Pearson squared correlation, $r^2$, between imputed genotype dosages and ground truth genotypes as a function of minor allele frequency, MAF, as estimated from the reference panel containing 1,100 present-day individuals ('1100 PD'). Each plot represents imputation accuracy for a sample size of 10 individuals with the same ancestry and age, which increases across the plots from left to right and top to bottom.

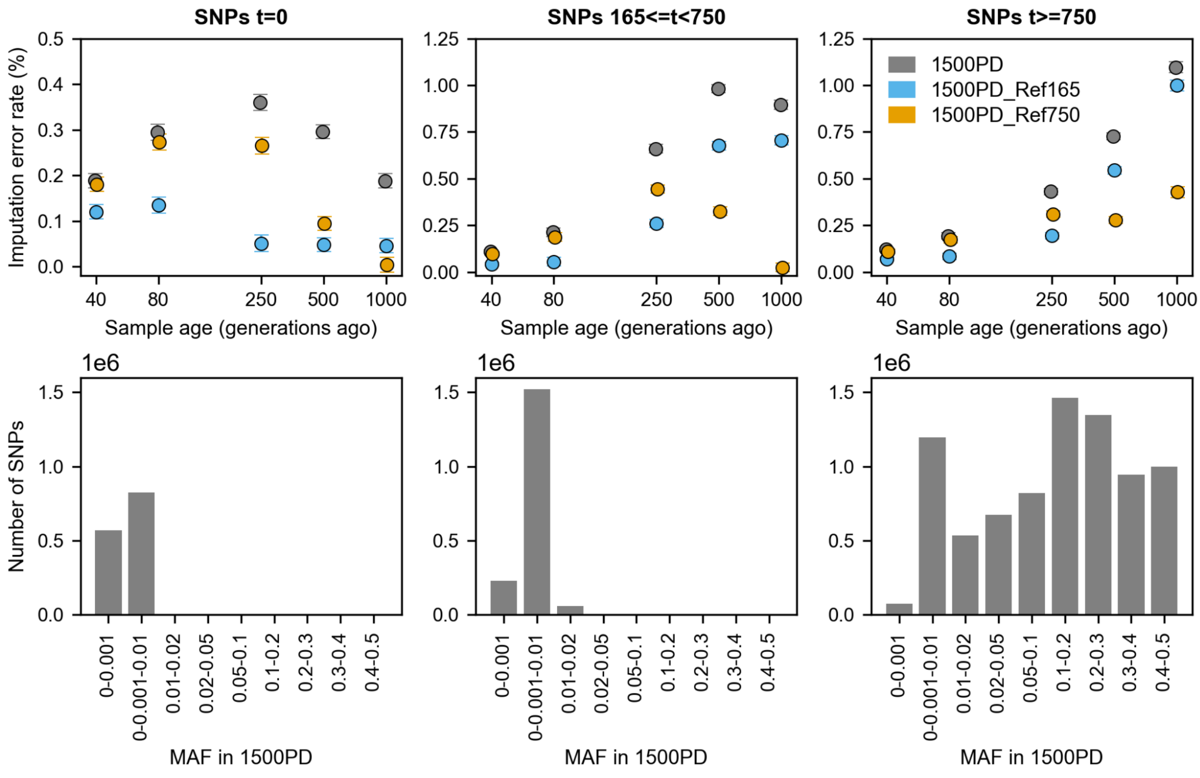

**Supplementary Fig. 13:** Imputation performance across sample ages and variant ages. We split variants based on which panel they appeared, from left to right: SNPs t=0 correspond to the variants that only existed at present time and not in Ref165 or Ref750; SNPs 165≤t<750 are variants that are found in Ref165 but not in Ref750; SNPs t≥750 are variants that were already present in Ref750. Top row: mean imputation error rate was calculated taking into account all sites when using different reference panels, namely, a reference panel with 1,500 present-day genomes ('1500PD'), and two panels based on '1500PD' to which we added 500 genomes from Ref165 ('1500PD_Ref165') or from Ref750 ('1500PD_Ref750'). The mean imputation error rate was calculated across ten individuals from each time point and we represent two times the standard error. Bottom row: number of SNPs per minor allele frequency (MAF) bin as estimated in the '1500PD' panel.

**Simulations of ancient individuals with distinct demographic histories**

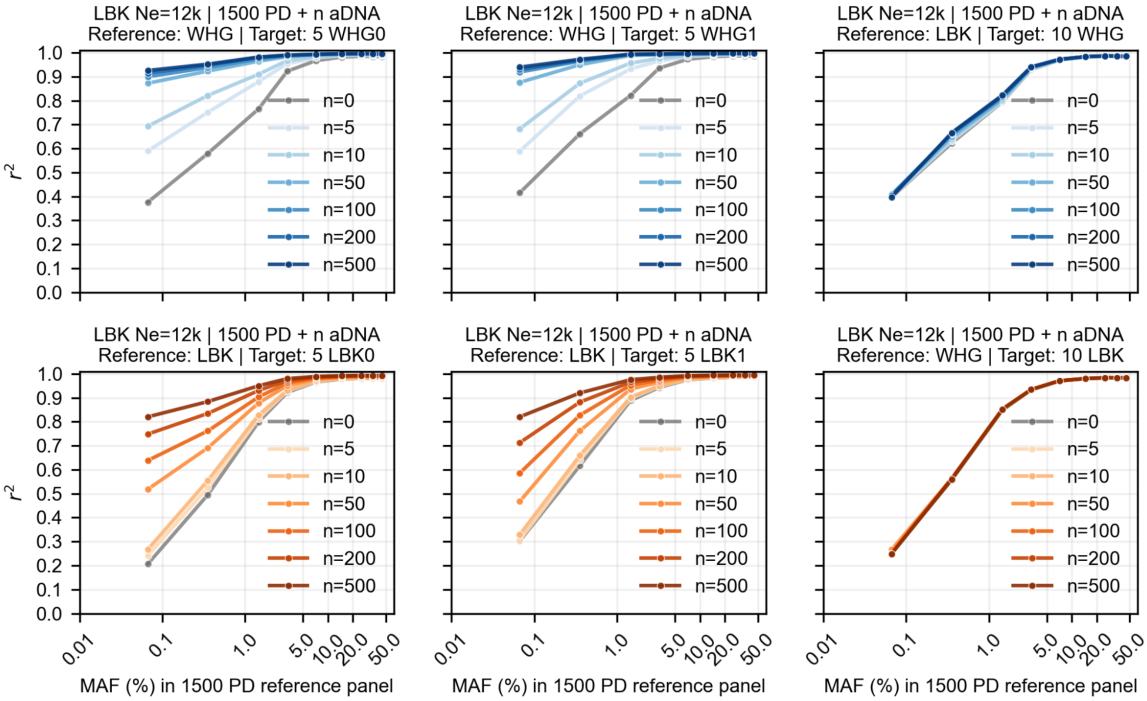


**Supplementary Fig. 14:** Effect of adding ancient haplotypes to the imputation reference panel when Ne of LBK-like populations equals 12,000. The different plots depict aggregate Pearson squared correlation, $r^2$, between imputed genotype dosages and ground truth genotypes as a function of minor allele frequency, MAF, as estimated from the reference panel containing 1,500 present-day individuals ('1500 PD'). Top row: imputation accuracy for 'WHG' target genomes; bottom row: imputation accuracy for 'LBK' target genomes. For the two first columns, we split imputation results for each target group into older and younger subpopulations (suffices 0 and 1, respectively) and the ancient reference haplotypes are well matched in ancestry; the third column shows imputation accuracy for the aggregate of the corresponding target population when the reference panel has different numbers of unmatched ancient reference haplotypes, i.e., imputation of 'WHG' using 'LBK' reference haplotypes and imputation of 'LBK' using 'WHG' reference haplotypes. Ne: effective population size; n: number of ancient genomes in the imputation reference panel.

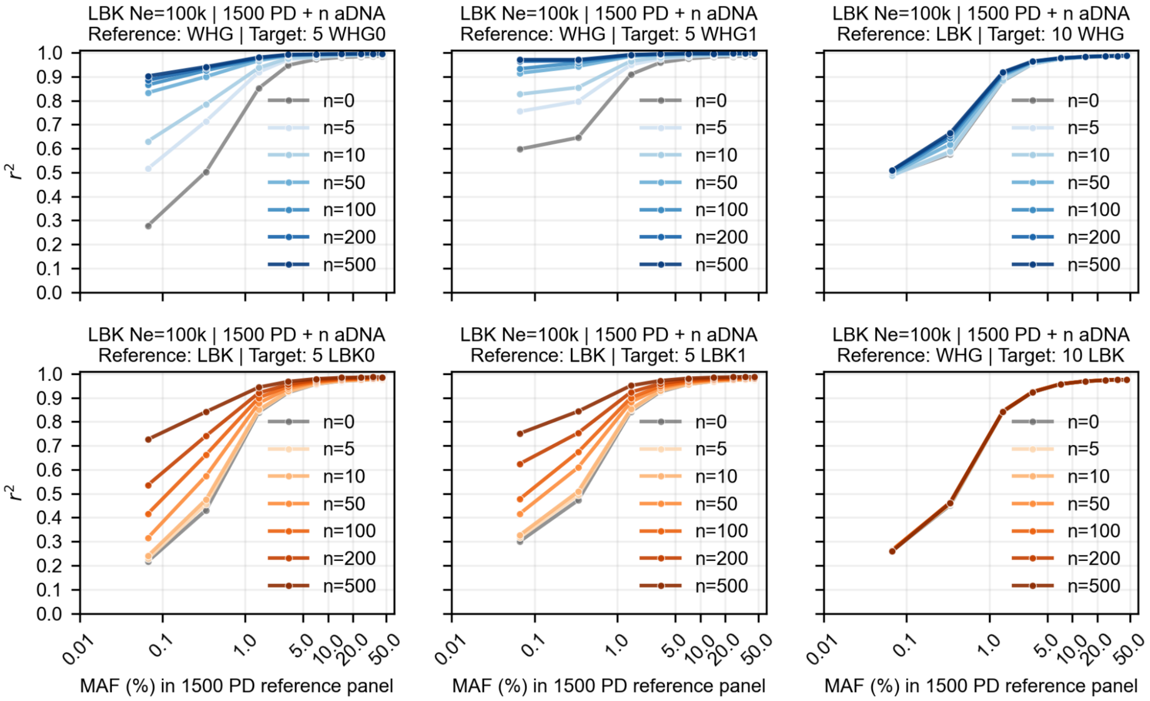


**Supplementary Fig. 15:** Effect of adding ancient haplotypes to the imputation reference panel when Ne of LBK-like populations equals 100,000. The different plots depict aggregate Pearson squared correlation, $r^2$, between imputed genotype dosages and ground truth genotypes as a function of minor allele frequency, MAF, as estimated from the reference panel containing 1,500 present-day individuals ('1500 PD'). Top row: imputation accuracy for 'WHG' target genomes; bottom row: imputation accuracy for 'LBK' target genomes. For the two first columns, we split imputation results for each target group into older and younger subpopulations (suffices 0 and 1, respectively) and the ancient reference haplotypes are well matched in ancestry; the third column shows imputation accuracy for the aggregate of the corresponding target population when the reference panel has different numbers of unmatched ancient reference haplotypes, i.e., imputation of 'WHG' using 'LBK' reference haplotypes and imputation of 'LBK' using 'WHG' reference haplotypes. Ne: effective population size; n: number of ancient genomes in the imputation reference panel.

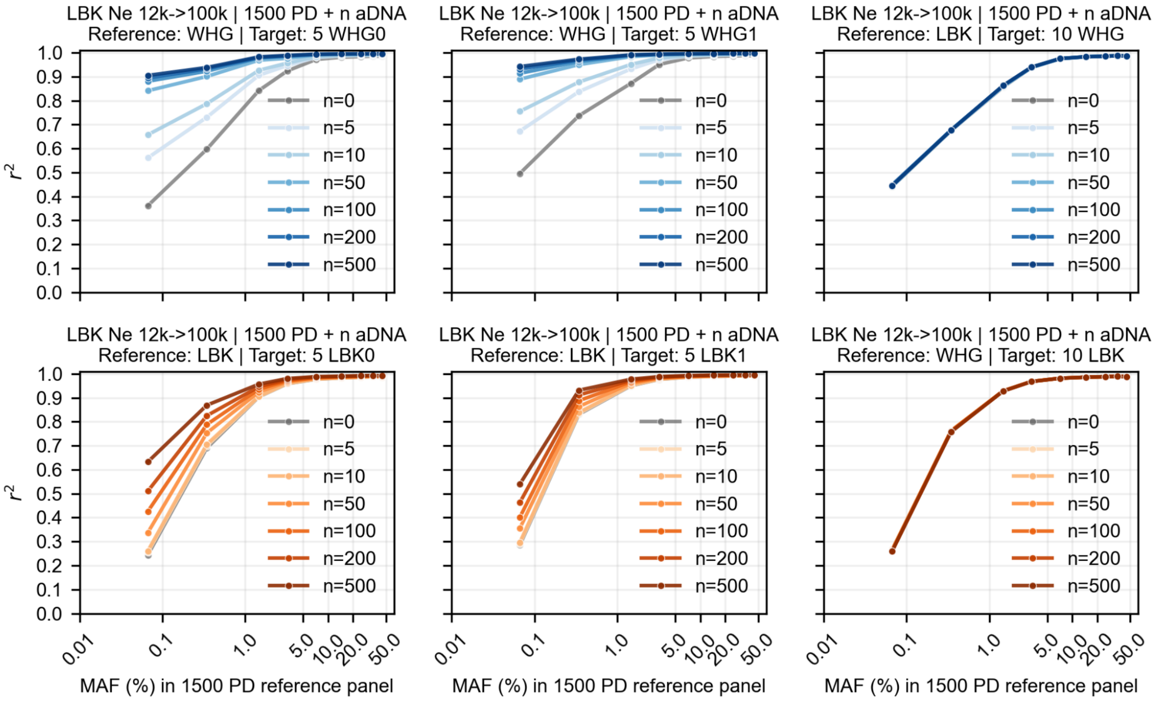


**Supplementary Fig. 16:** Effect of adding ancient haplotypes to the imputation reference panel when Ne of LBK-like populations expands from 12,000 to 100,000 at t=430 generations before present. The different plots depict aggregate Pearson squared correlation, $r^2$, between imputed genotype dosages and ground truth genotypes as a function of minor allele frequency, MAF, as estimated from the reference panel containing 1,500 present-day individuals ('1500 PD'). Top row: imputation accuracy for 'WHG' target genomes; bottom row: imputation accuracy for 'LBK' target genomes. For the two first columns, we split imputation results for each target group into older and younger subpopulations (suffices 0 and 1, respectively) and the ancient reference haplotypes are well matched in ancestry; the third column shows imputation accuracy for the aggregate of the corresponding target population when the reference panel has different numbers of unmatched ancient reference haplotypes, i.e., imputation of 'WHG' using 'LBK' reference haplotypes and imputation of 'LBK' using "Loschbour" reference haplotypes. Ne: effective population size; n: number of ancient genomes in the imputation reference panel.

### Supplementary Note 3 – An aDNA reference panel

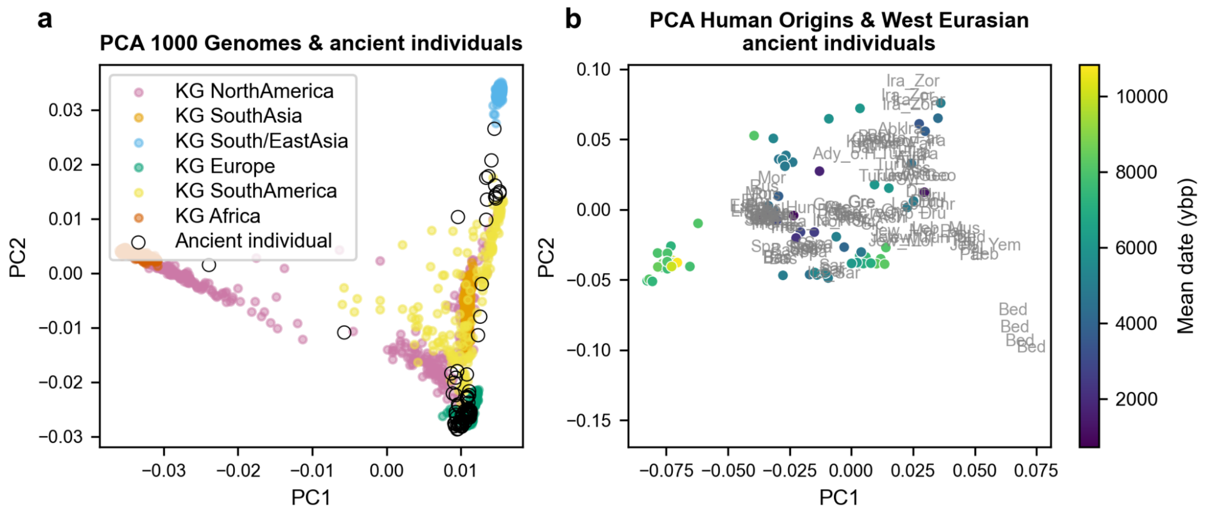


**Supplementary Fig. 17:** First two principal components from Principal component analysis (PCA) of high coverage ancient individuals in our dataset. Principal components (PC) were calculated using whole-genome data from present-day individuals and ancient genetic data was projected onto these (smartpca, lsqproject). a. 1000 Genomes individuals are colored by region (full circles) and ancient individuals coordinates are shown by black empty circles. b. Location in the first two PCs of West Eurasian Human Origins individuals are indicated by the first three letters of their labels and we are only plotting one for every six individuals to ease visualization; the PCs coordinates for 92 ancient individuals are represented by full circles colored by their corresponding mean date in years before present (ybp).

### Supplementary Note 4 – Testing genotype calling approaches for aDNA

Using the gargammel (Renaud et al. 2016) simulation of an ancient genome (chr1, 20x, based on individual genome NA20502 (TSI) from the 1000 Genomes dataset (“A Global Reference for Human Genetic Variation” 2015)), we tested different genotype calling approaches. We assessed i) how accurate the different approaches are (error rate) and ii) how many SNPs we can recover from each approach. Note that we are restricting our tests to known polymorphisms from the 1000 Genomes panel.

We tested the following calling approaches:

1. Bcftools basic (base and mapping quality)
2. Bcftools 5 filters (base quality, mapping quality, 1000 Genomes accessible mask, remove sites in repeat regions, QUAL>30, remove depth outliers (Moreno-Mayar et al. 2018))
3. bamRefine (Koptekin et al. 2025) followed by “bcftools basic”
4. bamRefine + bcftools + DP outliers + QUAL
5. bamRefine + bcftools + imputation

**3.1. Methods**

**3.1.2. Ancient reads simulation and mapping**

Using gargammel, we simulated aDNA-like reads using as a template the hg19 reference genome and one individual in 1000 Genomes (TSI, NA20502, phased data) without contamination (*--comp 0,0,1*) and with 20x coverage (*-c 20*). To introduce deamination at the ends of the reads, we used the same parameters as in (Koptekin et al. 2025) *-damage 0.024,0.36,0.0097,0.55*. For the fragment length distribution, we specified a log-normal distribution with parameters *--loc 4.106487474 --scale 0.358874723*, while restricting the fragments’ sizes to between 30 bp and 150 bp.

This simulation output paired-end reads that we subsequently mapped. First we removed adapters with AdapterRemoval v2.3.4, and then we mapped the reads with bwa v0.7.18 (*bwa aln -l 1024 -n 0.01 -o 2 -t 8 fasta reads.collapsed.gz*).

To verify whether the resulting genome had the intended characteristics (a 20x deaminated version of NA20502), we estimated postmortem damage rate with PMDtools (Skoglund et al. 2014), calculated genome coverage with samtools coverage and verified similarity with the original genotype calls after calling with bcftools (see Results). The simulated reads have high PMD rate at the ends of the reads (~35%) and show the typical pattern for a double-stranded library of C-to-T and G-to-A at the 5’ and 3’ ends, respectively (**Supplementary Fig. 17**). Furthermore, the resulting genome coverage is 19.67x and there is a high similarity to the original genome at the variant sites (~98%, see **Supplementary Fig. 18**).

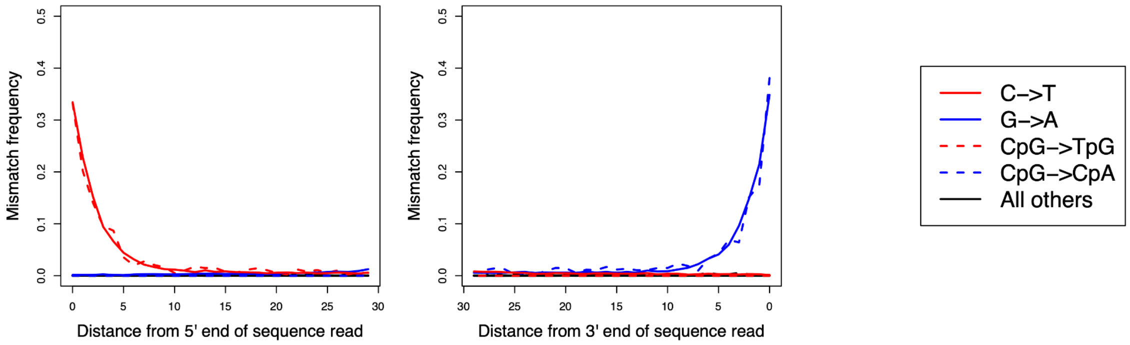


**Supplementary Fig. 18:** Deamination pattern at the 5’ and 3’ ends of the reads. Plot output by PMDtools.

##### 3.1.3. Genotype calling with bcftools

###### 3.1.3.1. “bcftools0”

We used the following command to call genotypes at the bi-allelic SNPs from the 1000 Genomes dataset:
*bcftools mpileup -f fasta -I -E -a 'FORMAT/DP' -q 30 -Q 20 --ignore-RG -T vcf -r 1 bam | bcftools call -Aim -C alleles -T tsv -Oz -o out*

###### 3.1.3.2. “bcftools5Filt”

For this callset, we applied some additional quality control filters: restricting variants to the 1000 Genomes accessible genome mask, removing positions in known repeat regions, removing depth outliers (minimum depth: ⅓ of the mean coverage; maximum depth: 2x mean coverage), and keeping only genotypes with QUAL>30 (Moreno-Mayar et al. 2018). We will refer to these two last filters as DP and QUAL, respectively.

##### 3.1.4. Applying bamRefine

We used bamRefine to mask C<->T and G<->A polymorphisms from the 1000 Genomes panel at the first L positions at the 5’ and 3’ ends, respectively. We used the same value for L (3, 5, 8 bp) at both reads’ ends.

##### 3.1.5. Imputation

We performed imputation using two different versions of the 1000 Genomes panel:

1. 1000 Genomes without TSI individuals (n=2397)
2. 1000 Genomes with only African and Asian populations (n=1497)

The first panel represents a case where the target genome is relatively well represented by the reference panel, for which we only removed individuals from the same population as the target (TSI), while the second mimics poorer genetic representation in the reference panel, with all genomes that could have European ancestry were removed. The latter panel aims to mimic situations in which the genetic diversity of ancient individual samples is not captured by present-day reference panels.

We used GLIMPSE2 (Rubinacci et al. 2023) for imputation and followed the steps from its tutorial (https://odelaneau.github.io/GLIMPSE/docs/tutorials). We used the same sets of chunks for the two imputation rounds. These chunks were calculated with GLIMPSE2_chunk on reference panel 1 (81 chunks). For the imputation step, we used genotype likelihoods as input as obtained from bcftools (see command in **1.1.3.1.**).

##### 3.1.6. Comparing genotype calls against the ground truth

To estimate error rates, for the files including genotype likelihoods (encoded by the FORMAT/PL field), we defined the resulting genotypes as the most likely genotype as defined by the PL field. For the imputed calls, the resulting genotype is the one with the highest genotype posterior (FORMAT/GP field). The ground truth dataset was directly extracted from the 1000 Genomes panel.

#### 3.2. Results

##### 3.2.1. Bcftools and bamRefine

As expected, calling genotypes with bcftools with base and mapping quality filters and no other quality control step led to the highest error rate, 1.62% (56,545 errors). Transition polymorphisms (impacted by PMD) were the most affected with an error rate of 2.19% against 0.28% for transversions (**Supplementary Table 8, Supplementary Fig. 18**). We found that using bamRefine to mask transitions potentially deaminated within three base pairs from the ends of the reads reduced the overall error rate to 0.73% (25,610 errors). In particular, the transitions error rate reduced to 0.94%, while the transversion error rates remained approximately the same (0.25%). The application of more stringent masking, that is, within a larger distance from the ends of the reads, led to further reductions in error rate. When we applied further quality control steps to calls generated with bamrefine followed by bcftools, we obtained even lower error rates, between 0.05% and 0.06% (2,027 and 1,610 errors for 3 bp and 8 bp, respectively). The bcftools 5 filters approach, which has been used in the literature in several studies (e.g., (Moreno-Mayar et al. 2018; Sousa da Mota et al. 2023)), produced one of the lowest error rates: 0.09% (2,199 errors), but it also yielded the highest missingness rate (32.75%). This approach was also particularly effective in reducing the error rate at transition polymorphisms with an error rate of only 0.12%, while the transversions error rate was still considerably lower (0.03%).

To summarize, the bamrefine + DP + QUAL approach led to the most accurate calls for almost all tested sites (bcftools 5 filters had lower error rate at heterozygous sites), while keeping the majority of sites with only around 10% of missing sites.

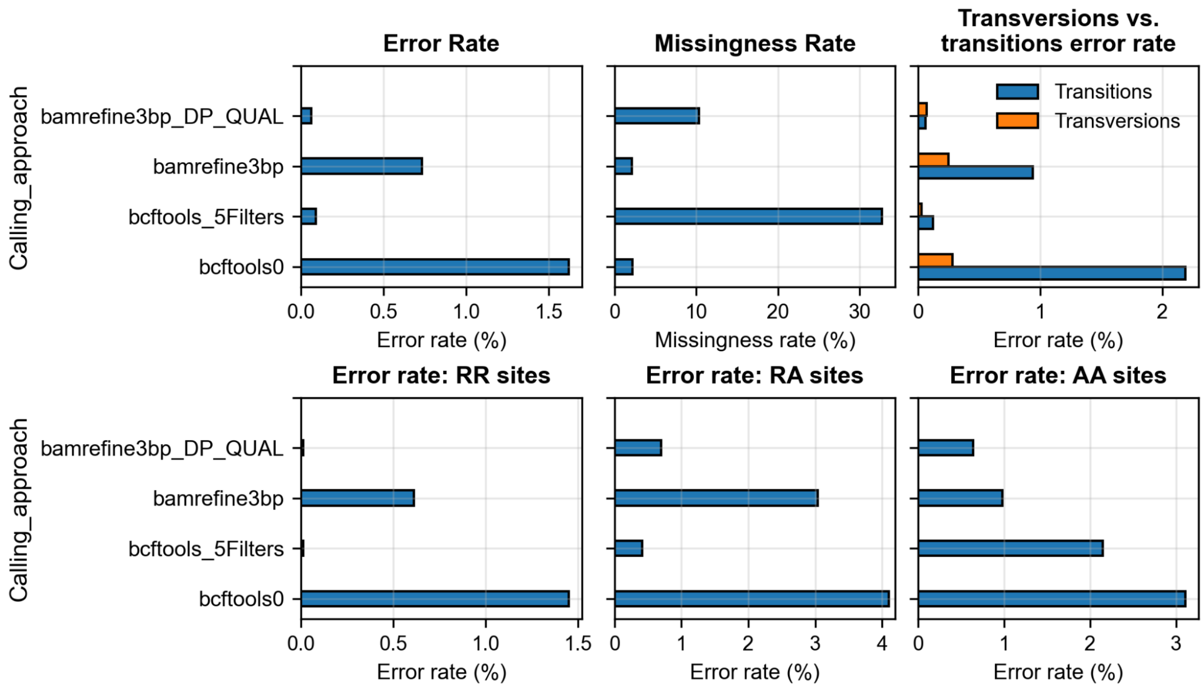


**Supplementary Fig. 19:** Assessment of different genotype calling approaches. From left to right and top to bottom: overall error rate (number of errors divided by the number of called sites), missingness rate (number of called sites divided by the number of biallelic SNPs for ground truth dataset), error rate for transversions vs. transitions, error rate at homozygous reference allele SNPs (RR), error rate at heterozygous sites (RA), and error rate at homozygous alternative allele SNPs (AA).

###### 3.2.2. Effect of using imputation to refine genotype calls

We also tested whether using imputation to refine genotype calls can lead to more accurate calls. We found that imputing after genotype calling had similar error rates as the bamRefine + DP + QUAL approach when bamRefine had been previously applied and the target ancestry was relatively well represented in the imputation reference panel (**Supplementary Fig. 19**, **Supplementary Table 8**). The overall error rate of bamRefine3bp followed by imputation with 1000 Genomes without TSI was 0.07% (bamRefine3bp + DP + QUAL had an error rate of 0.06%). As before, in both imputation experiments the application of bamRefine to remove deaminated sites led to lower error rates. Overall error rates decreased from 0.08% to 0.06% and from 0.12% to 0.09% for the reference panel without TSI and the reference panel containing only African and Asian genomes, respectively. When we discriminate the variants by the number of reference and alternative alleles, we found that alternative allele homozygous (AA) SNPs were better called when using imputation regardless of the panel and bamRefine usage, while calls at RR sites are more accurate with the bamRefine3bp + DP + QUAL approach. The lowest error rates were obtained when filtering imputed calls by their maximum genotype posterior (GP≥0.90): 0.04% and 0.05% for panels 1 and 2, respectively, and bamRefine 3 bp.

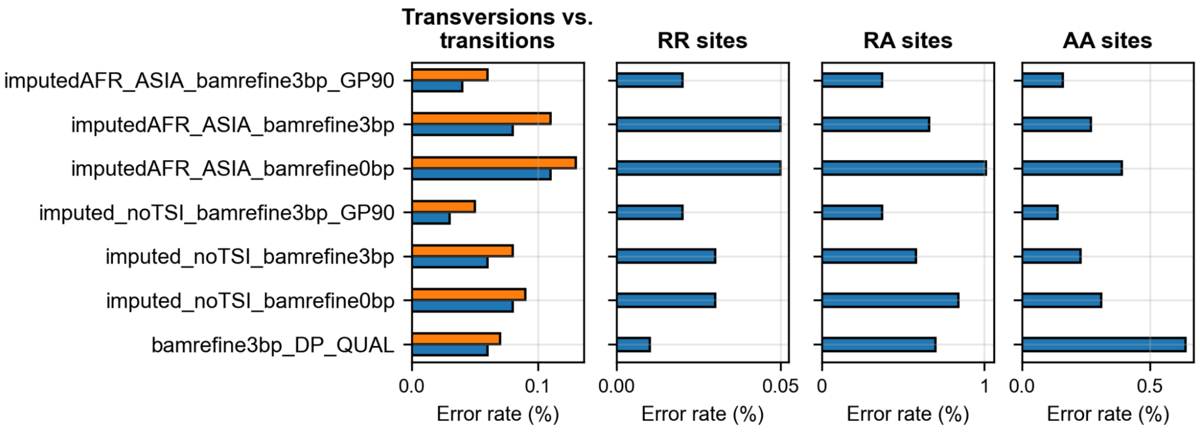


**Supplementary Fig. 20:** Effect of using imputation for genotype calling refinement on error rate. This figure includes results using two different imputation reference panels: 1000 Genomes without TSI (labeled as “impute_noTSI”) and 1000 Genomes including only populations from Africa and Asia (labeled as “AFR_ASIA”). From left to right, transversions vs. transitions error rates, error rates at homozygous reference allele SNPs (RR), error rate at heterozygous sites (RA), and error rate at homozygous alternative allele SNPs (AA).

##### 3.3. Conclusions

Here, we tested different genotype calling approaches anchored on bcftools. The bamRefine + DP + QUAL arose as a balanced approach that combines high accuracy and relatively low missingness. We also found that imputing after applying bamRefine to mask potentially deaminated variants and bcftools to generate genotype likelihoods leads to accurate genotypes. That is particularly the case when the imputation reference panel is well matched with the target in terms of genetic ancestry and after filtering for genotype posterior on the imputed calls.

We did not, however, test genotype callers specifically designed for ancient genomes, such as snpAD (Prüfer 2018) and ATLAS (Link et al. 2017), that can potentially produce equally accurate results.

### Supplementary Note 5 – Assessing phasing switch error rate

Phasing errors in an imputation reference panel may contribute to reduced imputation accuracy. To understand the magnitude of this problem, we estimated switch error rates using:

1. Simulated reads with deamination (gargammel (Renaud et al. 2016), 20x), based on an existing genome from 1000 Genomes (NA20502, see **Supplementary Note 4**)
2. High quality family data (aDNA) from the Afanasievo culture (dated to between ~4400 and ~4700 years before present): mother (I3388, 10x), father (I3950, 32x) and two children (I6714 and I3949, 25x and 30x, respectively) (Wohns et al. 2022)

**5.1. Methods**

**5.1.1. Phasing an aDNA family**

For the first case, we started by generating a ground truth dataset that we generated by phasing the genomes using pedigree information. We used as input data the genotype calls obtained by applying bamRefine (3 bp) (Koptekin et al. 2025), bcftools call with mapping quality 30 and base quality 20, and with depth and QUAL filters (see Methods, main text). We used SHAPEIT5 phase_common (Hofmeister et al. 2023) to phase the four relatives using family information and a reference panel (1000 Genomes).

We then phased each of the relatives separately using SHAPEIT5 phase_common and 1000 Genomes as the reference panel. We had two types of input data: bamrefine3p + bcftools + DP + QUAL and bamrefine3p + bcftools + imputation.

**5.1.2. Phasing a simulated genome**

As before, we used SHAPEIT5 to phase the calls for the simulated ancient genome as obtained from bamrefine3p + bcftools + DP + QUAL and bamrefine3p + bcftools + imputation. As in the **Supplementary Note 4**, we used two different versions of the 1000 Genomes panel so as to mimic a situation where the phasing/imputation reference panel does not match the genetic ancestry of the target genome well: i) 1000 Genomes without TSI individuals and ii) 1000 Genomes with only African and Asian genomes.

**5.1.3. Estimating switch error rates**

We used SHAPEIT5 switch to estimate switch error rates (SER) for both cases. For the simulated ancient genome, we used as ground truth the phased haplotypes for individual NA20502 as provided in 1000 Genomes.

**5.2. Results**

When phasing the simulated ancient genome, we found that using a reference panel that better represents the target ancestry, in this case, European, led to lower phasing errors (0.75-0.78% vs. 0.85-0.91%, **Supplementary Table 9**). We also found that applying imputation after genotype calling yielded lower SER, which was particularly evident for the real ancient family data. SER dropped from 0.81% (I3949) and 0.90% (I6714) to around 0.50% for both genomes.

**Supplementary Table 9:** This table contains SER (switch error rates) across different phasing experiments. 1KG: 1000 Genomes panel; 1KG_noTSI: 1000 Genomes panel without TSI (Toscani in Italy) genomes; 1KG_AFR_ASIA: 1000 Genomes panel without European or American genomes (only African and Asian).

| Sample ID | Sample type | Ground truth | Imputed target (yes/no) | Phasing reference panel | # Switch errors | # Sites | SER (%) |
|---|---|---|---|---|---|---|---|
| NA20502 | Simulated reads from real data | 1KG | No | 1KG_noTSI | 1090 | 140502 | 0.78 |
| NA20502 | Simulated reads from real data | 1KG | No | 1KG_AFR_ASIA | 1269 | 138900 | 0.91 |
| NA20502 | Simulated reads from real data | 1KG | Yes | 1KG_noTSI | 1163 | 155906 | 0.75 |
| NA20502 | Simulated reads from real data | 1KG | Yes | 1KG_AFR_ASIA | 1316 | 154017 | 0.85 |
| I3949.SG | Real family data (child) | Non-imputed, pedigree phased | No | 1KG | 614 | 75763 | 0.81 |
| I6714.SG | Real family data (child) | Non-imputed, pedigree phased | No | 1KG | 706 | 78597 | 0.90 |
| I3949.SG | Real family data (child) | Non-imputed, pedigree phased | Yes | 1KG | 398 | 76780 | 0.52 |
| I6714.SG | Real family data (child) | Non-imputed, pedigree phased | Yes | 1KG | 393 | 78845 | 0.50 |
| I3949.SG | Real family data (child) | Imputed, pedigree phased | Yes | 1KG | 784 | 156907 | 0.50 |
| I6714.SG | Real family data (child) | Imputed, pedigree phased | Yes | 1KG | 809 | 161267 | 0.50 |

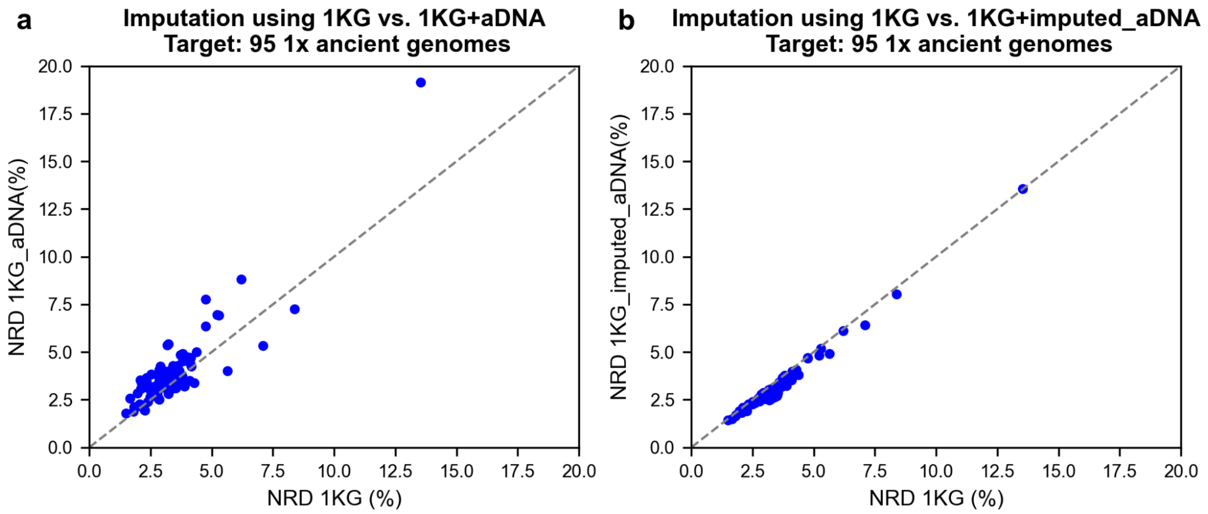


**Supplementary Fig. 21:** Imputation error rates when using different versions of the 1000 Genomes panel (1KG). Both plots depict non-reference discordance (NRD) for imputed calls of 95 ancient genomes downsampled to 1x. a. NRD comparison between imputation performed using a reference panel consisting of the 1KG alone (x-axis) and a merge of 1KG and 94 high-quality, non-imputed ancient genomes (y-axis). b. Effect on NRD when the high-quality ancient genomes were imputed before merging with 1KG.

##### 5.3. Conclusions

In our assessment of phasing errors, the two test cases have ancestries that are captured by present-day European populations that tend to be well represented by imputation/phasing reference panels. However, the assembled aDNA reference panel contains individuals from around the world with a diversity of genetic histories. To understand the impact of reference panel representation, we phased the simulated ancient genome using a reference panel without European populations (or genomes containing European ancestry via admixture). This experiment showed that poor genetic representation in the reference panel negatively impacts phasing performance. As such, the SER we estimated for the children from an Afanasievo family are likely to be a lower bound for SER for genomes of non-Western Eurasian ancestries.

Importantly, we obtained smaller phasing errors when the target had been imputed (~0.5% vs. 0.8-0.9% for the ancient family), which had been previously reported by (Medina-Tretmanis et al. 2024). We had shown in **Supplementary Note 4** that imputation can be used to refine genotype calls, yielding at minimum equally accurate calls, while allowing for a greater number of variants to be genotyped (at most, the number of variants in the reference panel). When adding real high coverage genomes to the 1000 Genomes, imputation errors were systematically equal or lower compared to using 1000 Genomes alone if the ancient reference genomes were previously imputed (**Supplementary Fig. 20**). Therefore, the improvement in phasing of the reference ancient haplotypes may be leading to improvements in imputation performance.

To improve imputation performance using ancient reference haplotypes, we will need to further improve phasing of those haplotypes. Large imputation reference panels have lower SER than what we found in our tests, for instance, the estimated SER for TOPMed with ~40K genomes varies between ~0.2% and ~0.3% (Browning et al. 2021). Increasing sample size is one of the most straightforward ways of reducing switch errors. As more high-quality ancient genomes are generated, we will probably be able to achieve such improvements.

### Supplementary Note 6 – Variant age and allele frequency effect on imputation

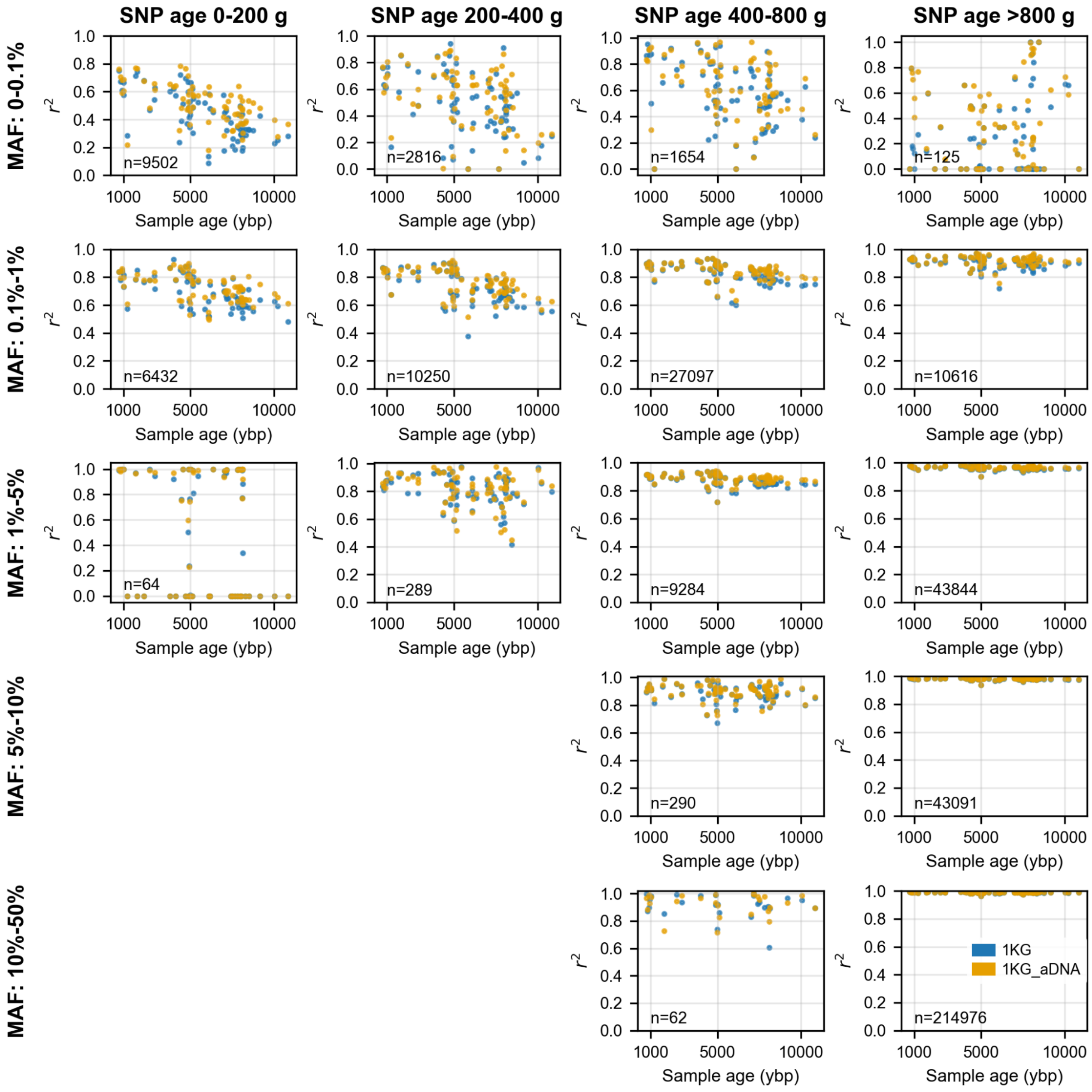


**Supplementary Fig. 22:** Imputation accuracy ($r^2$) for 1x West Eurasian genomes as a function of sample age in years before the present (ybp) for different SNP ages and minor allele frequencies (MAF). The data points in blue and yellow indicate results obtained when using 1KG (1000 Genomes) and 1KG_aDNA (1000 Genomes + ancient reference data), respectively. From left to right, $r^2$ estimated for variants binned by age in increasing fashion. From top to bottom, $r^2$ estimates start with very rare variants (0%-0.1% MAF) and end with very common variants (10%-50% MAF). We depict $r^2$ estimates calculated with a minimum of

51 SNPs. For each subplot, on the left lower corner, we represent the mean number of SNPs used in $r^2$ estimates across plotted data points. SNP ages have been calculated for CEU individuals (Utah residents with Northern and Western European ancestry) from the 1000 Genomes panel in (Speidel et al. 2019).
